# The wild grape genome sequence provides insights into the transition from dioecy to hermaphroditism during grape domestication

**DOI:** 10.1101/2020.01.07.897082

**Authors:** Hélène Badouin, Amandine Velt, François Gindraud, Timothée Flutre, Vincent Dumas, Sonia Vautrin, William Marande, Jonathan Corbi, Erika Sallet, Jérémy Ganofsky, Sylvain Santoni, Dominique Guyot, Eugenia Ricciardelli, Kristen Jepsen, Jos Käfer, Hélène Berges, Eric Duchêne, Franck Picard, Philippe Hugueney, Raquel Tavares, Roberto Bacilieri, Camille Rustenholz, Gabriel Marais

**Affiliations:** Université de Lyon, Université Lyon 1, CNRS, Laboratoire de Biométrie et Biologie Evolutive UMR 5558, F-69622 Villeurbanne, France; SVQV, Université de Strasbourg, INRA, Colmar, France; GQE–Le Moulon, INRA, Univ. Paris-Sud, CNRS, AgroParisTech, Univ. Paris-Saclay, 91190 Gif-sur-Yvette, France; INRA, Centre National de Ressources Génomiques Végétales, F-31326 Castanet-Tolosan, France; LIPM, Université de Toulouse, INRA, CNRS, Castanet-Tolosan, France; AGAP, Univ Montpellier, CIRAD, INRA, Montpellier SupAgro, Montpellier, France; PRABI, Université Lyon 1, F-69622 Villeurbanne, France; IGM Genomics Center, University of California, San Diego, La Jolla, CA.

**Author notes:** equivalent contribution as first authors (Hélène Badouin, Amandine Velt). equivalent contribution as senior authors (Roberto Bacilieri, Camille Rustenholz, Gabriel Marais).

## Abstract

Grapevine has a major economical and cultural importance since antiquity. A key step in domestication was the transition from separate sexes (dioecy) in wild *Vitis vinifera* ssp. *sylvestris* (*V. sylvestris*) to hermaphroditism in cultivated *Vitis vinifera* ssp. *vinifera*. While the grapevine sex locus is known to be small, its precise boundaries, gene content and the sex-determining genes are unknown. Here we obtained a high-quality *de novo* reference genome for *V. sylvestris* and whole-genome resequencing data of a cross. Studying SNP segregation patterns, gene content and expression in wild and cultivated accessions allowed us to build a model for sex determination in grapevine. In this model, up- and down-regulation of a cytokinin regulator is sufficient to cause female sterility and reversal to hermaphroditism, respectively. This study highlights the importance of neo-functionalization of Y alleles in sex determination and provides a resource for studying genetic diversity in *V. sylvestris* and the genomic processes of grapevine domestication.

---

Dioecy is rare in flowering plants (∼6%) but over-represented among crops (∼20%)^1^. In some cases, both wild and cultivated plants are dioecious (*e.g.* date palm, asparagus, persimmons). Other crops, such as grapevine, papaya, and strawberry, derive from dioecious progenitors and switched to hermaphroditism during domestication. The genes underlying this switch are currently not known in any crop. In *Vitis sylvestris*, wild females produce morphologically bisexual flowers, with retracted anthers that produce few and infertile pollen^2^, while male flowers undergo early ovule abortion^3^. Genetic analyses identified a 143-kb haplotype responsible for sex-determination in the grapevine reference genome^4, 5^. Two candidate genes for female sterility have been proposed, in particular *APRT3*^6^, a putative cytokinin regulator. However, the sequence of the sex locus was not available in *V. sylvestris* and no causative mutations have been identified. A genetic and evolutionary model to explain sex determination and switch to hermaphroditism in grapevine is currently lacking.

We sequenced a female individual of *V. sylvestris* using SMRT-sequencing (120X). Contigs were assembled with falcon-unzip^7^ and the grapevine reference genome^8^ (PN40024 – version 12X.2) was used to build pseudomolecules. We obtained a high-quality diploid assembly of 469 Mb with a contig N50 of 1.7 Mb, 98% of the gene content anchored on chromosomes and a BUSCO evaluation of 95% (Supplementary Table 1, Supplementary Figure 1), comparing favorably to other recently published plant genomes^9, 10^. We annotated 39,031 protein-coding genes on primary contigs.

To study sex determination, we localized the sex locus, which was fully included in the 5^th^ largest contig of the assembly. We also sequenced and assembled bacterial artificial chromosomes (BACs) covering the sex locus in a male of another *V. sylvestris* population and in *Vitis vinifera* cv. Cabernet-Sauvignon. To identify sex-linked polymorphisms in *V. sylvestris,* we produced a cross and resequenced in paired-end Illumina short reads the whole genomes of the parents and 10 offspring, yielding 43-74 millions of read pairs per individual (Supplementary Table 2). To overcome issues due to X-Y divergence, we used an iterative SNP-tolerant mapping procedure^11^, eventually mapping 98.2% of reads in average (mapping coverage by individual 14x-28x, Supplementary Table 3-6). Single nucleotide polymorphisms (SNPs) that segregate with sex in our cross were identified with an empirical approach and with SEX-DETector^++^, a new version of the probabilistic method SEX-DETector that identifies sex-linked genes from patterns of segregation in a cross from RNA-seq data^12^, which we developed here to analyse genomic data. As SNPs close to the sex locus might be genetically linked to the locus in a particular cross, we used public whole-genome resequencing data^13^ (Figure 1a-b, Supplementary Tables 7-8) to determine the boundaries of the sex locus. We searched for SNPs that were sex-linked in our cross and that were always heterozygous in males and homozygous in females in the validation dataset. We found that the X haplotype of the sex-locus spanned 111 kb on chromosome 2 (4,810,929 to 4,921,949 bp, Figure 1b-c, Supplementary Figures 2-5) and we obtained a final dataset of 1,865 XY SNPs (Supplementary Data File 1). To investigate changes associated with transition to hermaphroditism, we investigated the genotypes at the sex locus in publicly available DNA-seq data of 13 grapevine cultivars^14^ (Supplementary Table 9). Out of 13 cultivars, six harbored recombinant genotypes. The 5’ part of the sex locus was always either XH or HH and spanned 93 kb (4.810-4.903 Mb), H indicating the hermaphrodite haplotype, derived from the Y haplotype^5^. The remaining part of the sex locus (4.903-4.922Mb) was either XX or XH, but never HH (Figure 1b, Supplementary Table 10). The genes essential for the male phenotype should therefore be located in the 93 kb region of the sex locus.

**Figure 1:**
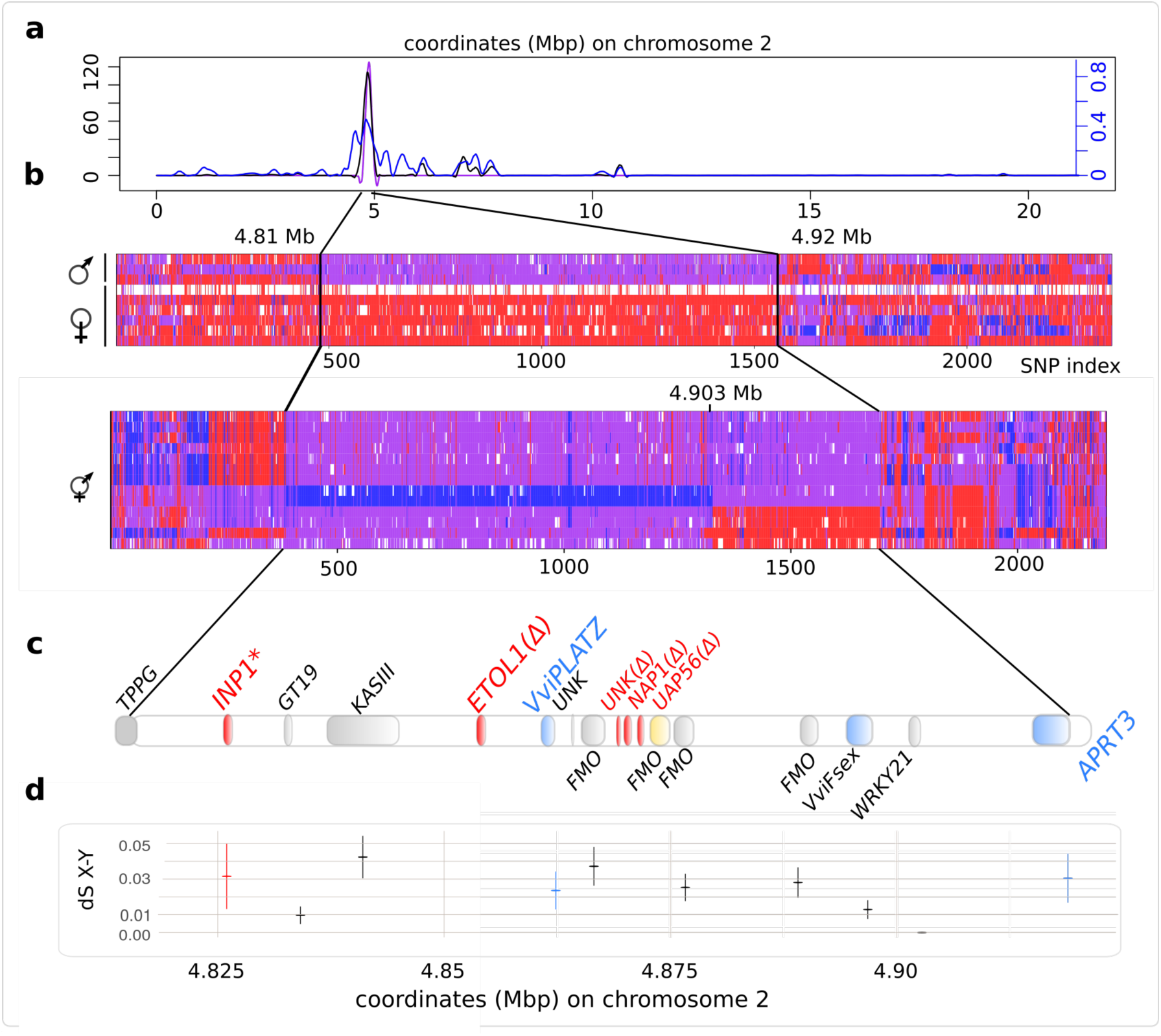
Limit, gene content and synonymous divergence in the sex locus of *V. sylvestris* and *V. vinifera*. **sa**, Detection of sex-linked SNPs on chromosome 2. left y axis: adjusted SNP number in 10kb windows, black curve: XY single nucleotide polymorphisms detected by SEX-DETector++ in a cross of *V. sylvestris*. Purple curve: candidate XY SNPs that show heterozygosity in males and homozygosity in females in a validation dataset of public whole-genome resequencing is drawn in purple; right y-axis (blue curve): adjusted mean posterior probability of being XY for SNPs in 10kb windows. **b,** Genotype of nine *V. sylvestris* individuals (top) and thirteen grapevine cultivars (bottom) in a validation dataset of public whole-genome resequencing, at locations of candidates XY SNPs. Red, purple and blue, and white traits represent XX, XY, YY and missing genotypes, respectively. The black lines highlight the limits of shared XY SNPs between the cross resequenced in the present study and the validation dataset. **c,** Gene content and annotation and in the sex locus (approximate position). Genes highlighted in red are absent (Δ) or possess a frameshift deletion in the X haplotype. Genes highlighted in blue are induced in males. The yellow gene is absent in the Y haplotype. **d,** synonymous divergence between X and Y allele in the sex locus (+/− standart error), reflecting the age of recombination suppression. dS was only computed for genes present in both haplotypes.

Next, we investigated the age of the sex locus, and the presence of deletions and insertions of transposable elements in the Y haplotype that may indicate degeneration. We calculated the synonymous divergence (dS) between X and Y coding DNA sequences (CDS) of XY gene pairs (Figure 1c) as a proxy of the age of the system (Figure 1d). The two genes at the opposite limits of the non-recombining region had a dS of 0.03, suggesting that suppression of recombination occurred very recently and in a single step in *V. sylvestris*. The maximum dS was 0.0424, which is to our knowledge lower than any of the other systems dated in plants (*Mercurialis annuua* has a maximum dS of 0.05^15^). To identify deletions in the Y haplotype, we compared the X haplotype and two BAC contigs covering the Y haplotype (Supplementary Figures 2-5). We also measured the mean mapping coverage in our resequencing dataset, searching for two-fold reductions in males. This revealed eight regions spanning 500 to 5,500 bp and affecting one predicted gene (Supplementary Figure 5). We also detected insertions of transposable elements in the Y haplotype (Supplementary Figures 2-5, Supplementary Data File 2). This suggests that the Y haplotype of *V. sylvestris* is already degenerating despite recombination suppression being very recent.

Theoretical work predicts that XY sex chromosomes should combine a recessive male-sterility mutation (X) and a dominant female-sterility mutation^16^ (Y). To identify candidates, we compared the gene content of X, Y and H haplotypes (Figure 2a) and searched for presence/absence patterns and loss-of-function mutations (Figure 2b). We also mapped a public RNA-seq dataset of males, females and hermaphrodites flower buds of *V. sylvestris* at four developmental stages^13^ (Supplementary Table 11), measuring the total (Figure 3a) and the allele-specific (X, Y and H, Figure 3b) expression of transcripts.

**Figure 2:**
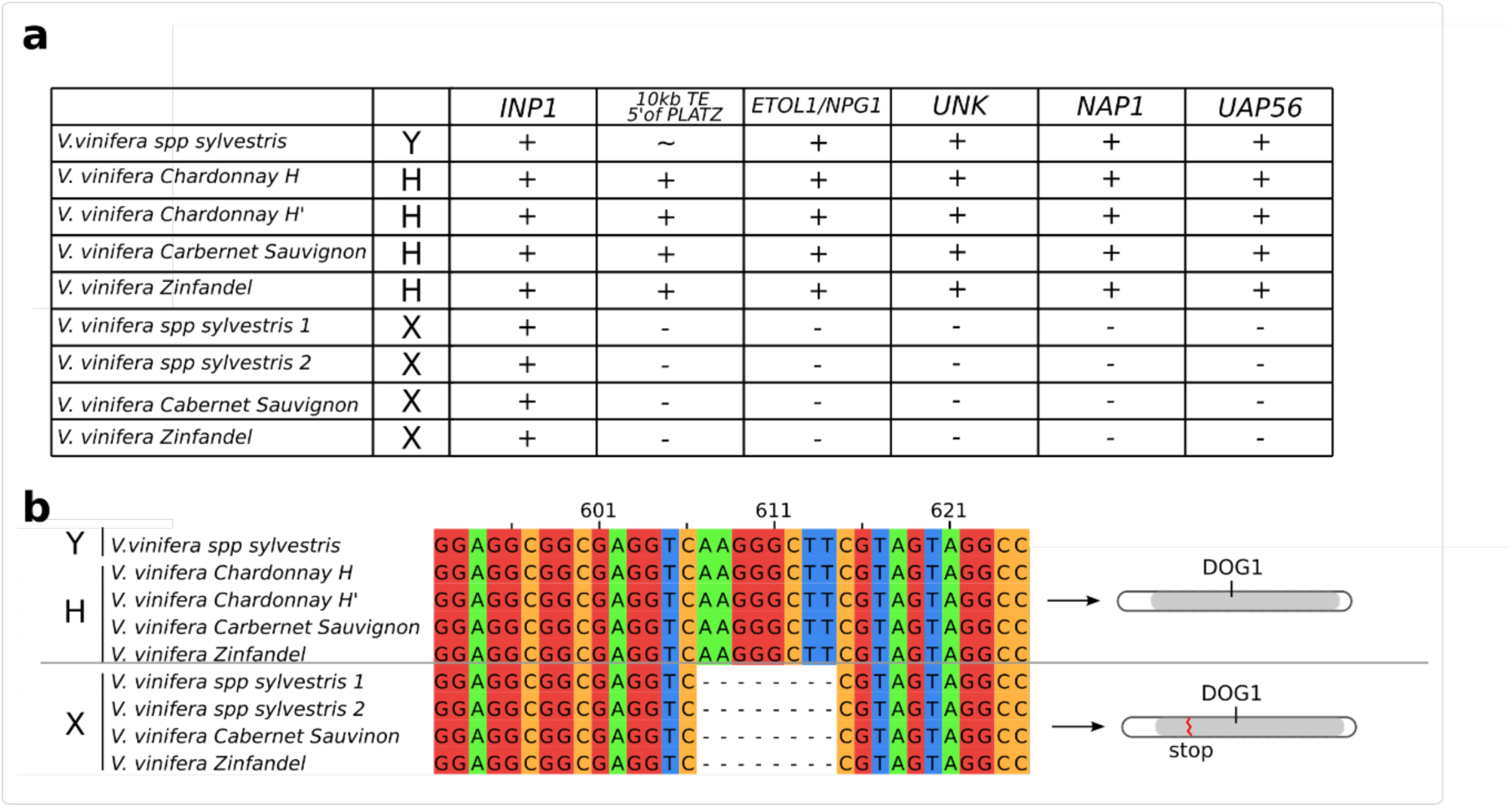
Presence-absence patterns and frameshifts mutations in X, Y and H haplotypes. **a**, Presence-absence patterns of five genes absent or truncated in the X haplotype of *V. sylvestris*, and of a 10kb repeated element 1.2 kb uptream of a gene of the PLATZ family of transcription factors. The tild indicates that the estimated length of the gap in the Y haplotype inferred is consitent with the presence of the element, but is has not been sequenced yet. **b,** An 8-bp deletion in exon 2 of the *INP1* gene is shared by X haplotypes and absent in Y or H haplotypes. It results in a premature stop codon, leading to a shorter protein. The DOG protein domain, involved in DNA binding, is truncated.

First, we investigated the presence of X recessive mutations possibly to causing male sterility. We found that the gene at one limit of the sex locus, annotated as *INAPERTURATE* (*INP1)*^17^, showed a 8-bp deletion in exon2 in all female haplotypes, resulting in a premature stop codon and a truncated protein (Figure 2b). *INP1* loss-of-function mutant in *A. thaliana* lacks pollen apertures^18^, similarly to the pollen of female *V. sylvestris*^2^. We found that *INP1* was highly specific of mature flower buds of *V. vinifera*, consistent with a role in late pollen development^19^. It remains to be shown that the absence of apertures in grapevine female pollen is sufficient to cause sterility, as inaperturate sterile pollen evolved independently at least six times in Eudicots in association with dioecy^20^, but the *A. thaliania INP1* knock-out mutant is fertile^19^. We also found that four genes were absent from the X haplotype (Figures 1a-2b Supplementary Figure 2). They included a gene previously annotated^5^ as a short homologue of Ethylene-overproducer-like-1 (*ETOL1)*, but also showing homology to *NPG1*. *NPG1* is essential to pollen germination in *A. thaliana*^21^, and the ethylene pathway has been shown to be determinant in floral morph determination in *Cucumis melo*^22^. Three other genes were also absent from the X haplotype, namely a gene encoding a short peptide of unknown function, *NAP1* and *UAP56A*. *NAP1* encodes a nucleosome assembly protein in *A. thaliana*. Two homologues of *UAP56A,* a DEAD-box ATP-dependent RNA helicase, have been shown to regulate programmed cell death during tapetum development in *Oryza sativa*^23^, and disrupting them led to male sterility. None of these three genes were significantly expressed in the early developmental stages of the RNA-seq dataset that were produced to study ovule development (Supplementary Figure 6), likely because male sterility occurs later than female sterility in *V. sylvestris* flowers^13^. *INP1* and the four X-deleted genes were all present in at least one haplotype in cultivars. These results suggest that several recessive deletion mutations affecting genes involved in tapetum and pollen development may cause the male sterility syndrome (Figure 4).

**Figure 3:**
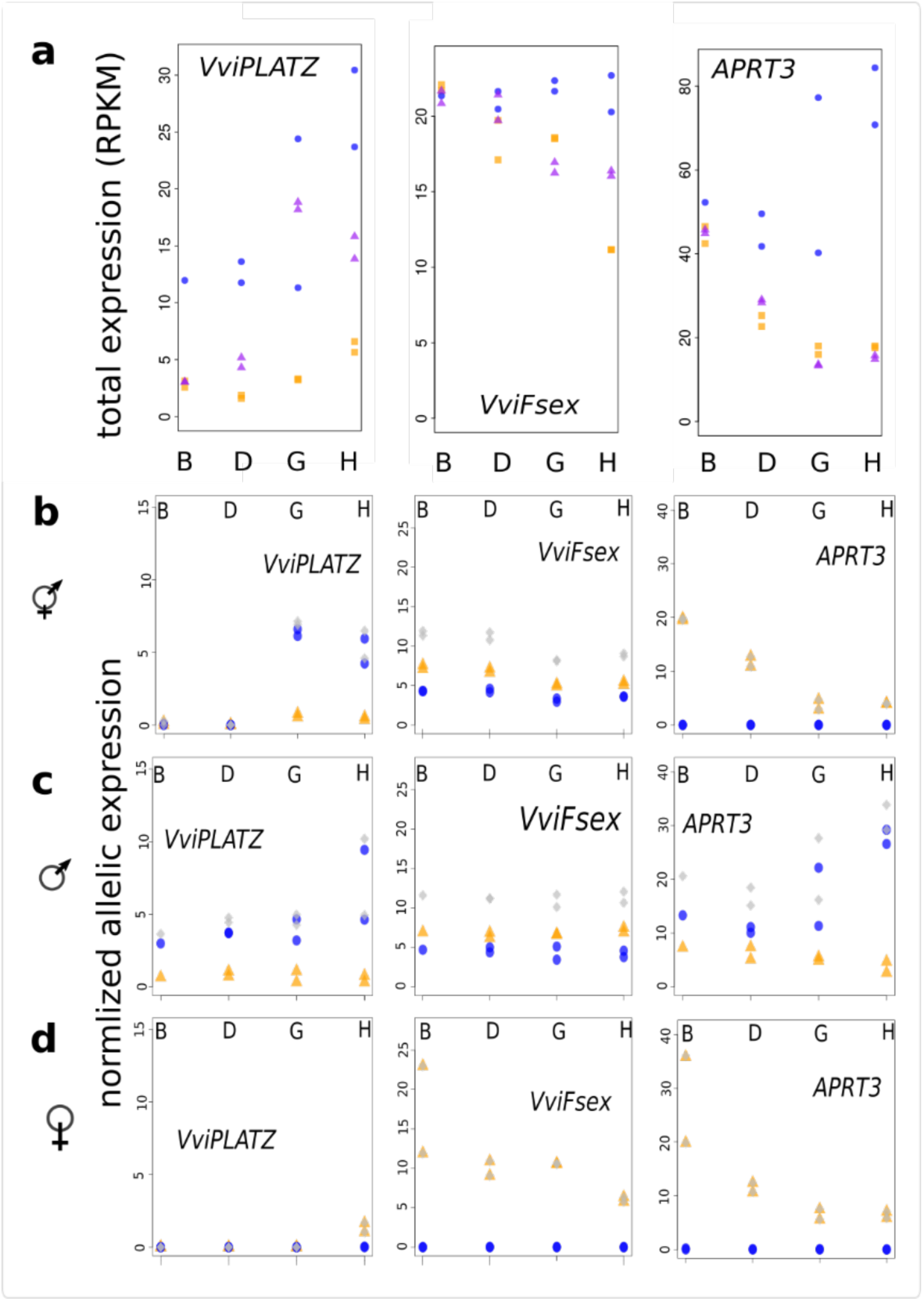
Total and allele-specific expression of female-sterility candidates during flower bud development. B, D, G and H represent early development stages of flower development sequenced in RNA-seq in Ramos et al^13^. **a,** total normalized expression (RPKM) of *VviPLATZ*, *VviFsex* and *APRT3* in males (blue circles), females (orange squares) and spontaneous hermaphrodites (purple triangles) of *V. sylvestris*. Each point represents a biological replicate. **b,** allele-specific normalized expression. Blue dots, orange triangles, and grey diamonds represent Y-specific, X-specific and summed expression, respectively. RNA-seq data were genotyped and the coverage of X and Y variants was extracted from vcf file and averaged by gene. Read counts were normalized by library.

**Figure 4:**
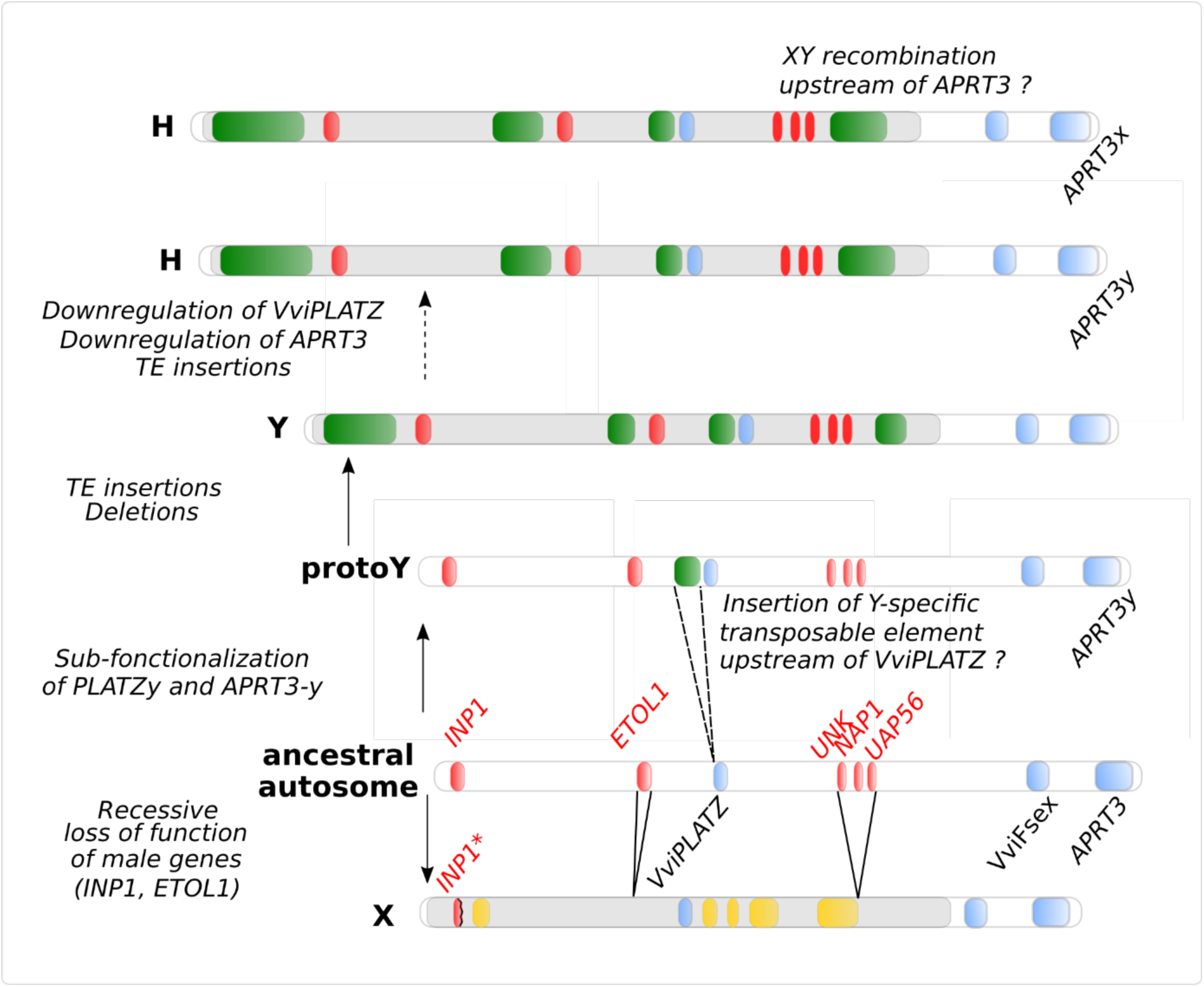
Evolutionary genomic scenario of the formation of the X and Y haplotypes and reversal to hermaphroditism. Red: genes partially or completely deleted in the X haplotype. Blue: genes differentially expressed between male, female and hermaphrodite during early flower bud development. On the schematic sex locus, yellow highlights represent X-hemizygote regions (*i.e.*, absent from the Y), green highlights represent transposable elements, red highlights male-sterility candidates and blue highlights female-sterility candidates. Grey highlight represents the part of the sex locus where homology between X and Y is reduced due to deletions or insertions.

Second, we searched for dominant mutations causing female sterility in the Y haplotype and for possible mechanisms of reversion to hermaphroditism. So far, sex-determining genes that have been identified in plants were Y-specific genes that arose through duplication^24–26^. The Y-specific genes described above (*INP1*, *ETOL1*, *UNK* and *UAP56*) are not expressed at the stage where ovule abortion occurs and are unlikely candidates for female sterility (Supplementary Figure 6). Therefore, we searched for differential expression of X and Y alleles in XY gene pairs. Three genes showed higher total expression in males than in females in the sex locus: *VviPLATZ*, *VviFSEX* and *APRT3* (Figure 3a). *VviPLATZ* and *APRT3* showed a similar expression pattern in males, with a two-fold induction during bud development. However, *APRT3* was also expressed in female while *VviPLATZ* expression was male-specific. Interestingly, only the Y alleles of *VviPLATZ* and *APRT3* (*VviPLATZy* and *APRT3y*) were induced in male buds (Figure 3b, Supplementary Figure 7). *VviFSEX* showed a decreasing versus stable expression in female and males, respectively, with equal expression of its X and Y alleles. The promoters of *APRT3x* and *APRT3y* were structurally similar, but a Y-specific transposable element may be present upstream of *VviPLATZy* (Supplementary Text), which could explain its up-regulation. *APRT3* is the ortholog of a gene encoding an enzyme of cytokinin elimination. Exogenous application of cytokinin is sufficient to restore female fertility in male *V. sylvestris*^27^. Previous *In situ* hybridisation work suggested that APRT3 was induced during male bud development and absent in female buds^6^. Here we found that *APRT3x* is expressed both in female and male buds, and is an ubiquitously expressed gene in *V. vinifera* (Supplementary Figure 8, Supplementary Table 12). Since all analysed cultivars possessed at least one *APRT3x* allele (recombinant *APRT3x/APRT3x* genotypes were observed but no *APRT3y/APRT3y*, Figure 1b), we hypothesize that *APRT3x* performs essential functions in *V. sylvestris* and in grapevine. *VviPLATZ* is a transcription factor of the PLATZ family. Interestingly, in the RNA-seq dataset, we found that *VviPLATZy* was expressed at a lower level in hermaphrodites than in males and that *APRT3* was not induced as in males (since it is RNA-seq data, the hermaphrodite genotype at *APRT3* is however unknown, Figure 3a). A tentative scenario for female sterility would be that *VviPLATZy* specifically activates *APRT3y*, leading to a decrease in cytokinin concentration in ovule and causing its abortion (Figure 4). Interestingly, a 10 kb repeated element is present 1.2 kb upstream of *APRT3* in all H haplotypes but absent from X haplotypes (Figure 2a), and might influence the expression of *APRT3*. From an evolutionary perspective, female sterility in grapevine would originate from neo-functionalization of the Y allele of an ubiquitous gene, the X allele conserving the ancestral function. Down-regulation of *VviPLATZy* and/or *APRT3y* may be sufficient to cause reversal to hermaphroditism (Figure 4, Supplementary Text). In addition (but not in a mutually exclusive way), transition to hermaphroditism may be caused by recombination upstream of the *APRT3* promoter leading to the *APRT3x/APRT3x* genotype that is observed in several cultivars (Figure 1b).

Following an extensive analysis of the *V. sylvestris* sex locus, we have proposed a model for sex determination in grapevine, based on a combination of deletion of X alleles and neo-functionalisation of Y alleles. In our model, the switch from dioecy to hermaphroditism was easy, as a single genomic change (possibly two) was needed. This is in agreement with recent studies^28^ suggesting that dioecy is rare in flowering plants because it has a high reversion rate to hermaphroditism. Future genetic and functional studies will allow dissecting the role of the pinpointed genes in different aspects of the male sterility syndrome, female sterility and reversal to female fertility in hermaphrodites.

## Data Availability

All raw reads, genome assembly and gene annotation are currently under submission.

## Code Availability

The source code of SEX-DETector++ is available at the url https://gitlab.in2p3.fr/sex-det-family/sex-detector-plusplus.

## Acknowledgements

This work was supported by ANR grants ANR-13-BSV6-0010 to PH and ANR-14-CE19-0021-01 to GABM., an INRA starter grant BAP2014_44–SELVI to TF and UMT Génovigne<u>®</u> grant to RB. This work benefitted from the computing facilities of the CC LBBE/PRABI. Whole genome PacBio was conducted at the IGM Genomics Center, University of California, San Diego, La Jolla, CA. RNA-seq and cross resequencing was performed by the GenomEast platform, a member of the “France Génomique”consortium (ANR-10-INBS-0009). Sylvestris C1-2 illumina resequencing was performed by the Etude du Polymorphisme des Génomes Végétaux (EPGV, INRA, Evry) facility. We thank Jérôme Gouzy for help with curation of the trinity dataset used for gene annotation and Aline Muyle for providing information on the code of SEX-DETector perl, Gisèle Butterlin and Angélique Ardiller for technical assistance and the SEAV experimental platform (Service d’expérimentation Agronomique et Viticole) for plant material maintenance at INRA Colmar. The authors would like to acknowledge support of GeT-PlaGe Genomic Platform (INRA-Toulouse, France, https://get.genotoul.fr) for providing PacBio BAC sequence data.

## Contributions

GABM, PH, CR, ED, RT and RB conceived the research. VD and ED obtained plant material and DNA for cross resequencing and genome sequencing. WM obtained DNA for whole genome PacBio sequencing. TF and RB obtained plant material and DNA for BAC sequencing, and with SS defined probes for BAC screening. SV and HBe performed BAC library production and screening, BAC sequencing and assembly. AV, CR and JC assembled the genome. RB and TF analysed BAC sequences and assigned them to sex haplotyes. SS sequenced the gaps between BAC contigs. WM, HBe, ER, KJ and CR coordinated the sequencing of the genome. ED and CR coordinated the sequencing of the transcriptome. GABM coordinated the sequencing of the cross. AV, CR and JC assembled the genome. JG and RT assembled the transcriptome. AV, CR and ES annotated the genome and the BAC sequences. FG, FP, DG, JK and GABM developed SEX-DETector++. HBa, AV and CR analysed RNA-seq data. HBa analysed WGS data and performed structural and molecular evolution analysis of the sex locus. HBa, CR and RB performed integrative analysis of sex determination. PH, RT, GABM and RB funded the study. CR, GABM and RB coordinated the project. HBa and GABM wrote the manuscript, with contributions from all authors.

## Guide to Supplementary Material

### Material and Methods

- Statistical analyses and data visualisation.
- Sequencing data
- Genome assembly, cleaning, polishing, anchoring and ordering.
- BAC generation and sequencing.
- Genome and BAC annotation.
- Development of SEX-DETector++.
- Characterization of the sex-linked region in the *V. sylvestris* genome.
- Integrative search for sex-determining genes in the sex locus.
- Supplementary References

### Supplementary figures

- Supplementary Figure 1: BUSCO analysis of the genome annotation.
- Supplementary Figure 2-4: Structural comparisons of X and Y, X and H and Y and H haplotypes, respectively.
- Supplementary Figure 5: Density of sex-linked SNPs in a cross between two *V. sylvestris* parents.
- Supplementary Figure 6: Total normalized gene expression in males, females and hermaphrodites of *V. sylvestris* for genes in the sex locus during flower bud development.
- Supplementary Figure 7: Allele-specific expression of X and Y alleles of XY genes pairs of *Vitis sylvestris* during the development of flower buds.
- Supplementary Figure 8: Distribution of organ-specificity expression of genes in *Vitis sylvestris*.

### Supplementary tables

- Supplementary Table 1: Genome assembly and anchoring statistics
- Supplementary Table 2: Summary statistics of resequencing data of a cross between two *V. sylvestris* parents
- Supplementary Table 3: Mode of the gaussian distribution of mapping coverage for each sample for the dataset of whole genome resequencing of a cross in *V. sylvestris*.
- Supplementary Table 4: Summary statistics of iterative SNP-tolerant mapping for the resequencing dataset of a cross in *V. sylvestris*.
- Supplementary Table 5: Summary statistics of missingness per sample for each iteration of SNP-calling in the dataset of whole-genome resequencing of a cross in *V. sylvestris*.
- Supplementary Table 6: Number of homozygous and heterozygous SNPs called at each iteration in the dataset of whole-genome resequencing of a cross in *V. sylvestris*.
- Supplementary Table 7: Whole-genome resequencing dataset of the ncbi short read archive that were mapped to the *Vitis sylvestris* genome.
- Supplementary Table 8: Statistics of mapping and SNP-calling of a public whole-genome resequencing dataset of the ncbi that was mapped to the *Vitis sylvestris* genome.
- Supplementary Table 9: Summary of the genotype of 13 cultivars in the sex locus on chromosome 2 inferred from whole-genome resequencing data.
- Supplementary Table 10: Predicted genes in the sex locus of *V. sylvestris*. Coordinates and geneID are indicated for the *V. sylvestris* reference genome.
- Supplementary Table 11: Mapping statistics of public RNAseq libraries of flower buds of female, male and hermaphrodite *V. sylvestris* against the *V. sylvestris* genome.
- Supplementary Table 12: Public RNAseq dataset of *V. vinifera* and *V. sylvestris* mapped to the *Vitis sylvestris* genome.

### Supplementary text

This text describes the results of BAC assembly and the attemps to characterize and to cover the 13kb gap on the male haplotypes.

### Supplementary data files

- Supplementary_DataFile1.xlsx: genomic location of sex-linked single nucleotide polymorphisms in the genome of *Vitis sylvestris*.
- Supplementary_DataFile2.xlsx: genomic location of repeated elements in the genome of *Vitis sylvestris*.
- egnep.Vvi.for_Sylvestris.cfg: configuration file of Eugene annotation.

## Supplementary Information

### Material and Methods

#### Statistical analyses and data visualisation

Unless stated otherwise, statistical analyses were carried out in R v3.4.4 (2018-03-15)^29^. Data visualisation were performed with R or with circos v0.69-6^30^. Adeneget^31^ was used to visualize SNP data in R.

#### Sequencing data

##### Whole genome sequencing (PacBio – long reads) of Sylvestris C1-2

The accession of Sylvestris C1-2 with plant code 8500.Col.C1-2, origin of the geographic location Sainte-Croix-en-Plaine, Haut-Rhin (68), France, was used for extraction of high-molecular-weight genomic DNA. The extraction was performed by the CNRGV (Centre National de Ressources Génomiques Végétales) – INRA Toulouse. They started with 1G of frozen material (−80°C), crushed it with liquid nitrogen and used the Genomic-tip 100/G kit (Qiagen) for the extraction. The SMRT library preparation and sequencing on PacBio RSII platform (P6-C4 chemistry) was done by the IGM Genomics Center at the University of California, San Diego, following the standard PacBio protocols. A total of 129X coverage was obtained in order to perform *de novo* genome assembly.

##### Whole genome sequencing (Illumina – short reads) of Sylvestris C1-2

DNA libraries with two different library size from leaves of Sylvestris C1-2 were prepared in order to perform 2*151 paired-end short reads sequencing (Illumina). One library had a library size of 740bp (860 bp with adapters) and the other library had a library size of 392bp (512bp with adapters). The two libraries were sequenced on the same flowcell lane. The libraries preparation and Illumina sequencing was performed by the EPGV (Etude du Polymorphisme des Génomes Végétaux) – INRA. A total of 338,109,086 reads was obtained (∼100X coverage) for all samples together, used to polish the Sylvestris C1-2 PacBio assembly.

##### RNA-sequencing of Sylvestris C1-2

RNA-sequencing was performed on six samples of the Sylvestris C1-2 accession. Three biological replicates of whole green berries and three biological replicates of whole mid-ripening berries were sequenced. The RNA-seq library preparation was performed with the TruSeq Stranded mRNA Library Prep Kit (Illumina) and was sequenced in paired-end (2*100b) on HiSeq4000 platform (Illumina technology). Library preparation and sequencing was performed by the GenomEast platform – Strasbourg. A total of 1,113,531,260 reads was obtained for all samples together, used to assemble a Sylvestris transcriptome and perform genes annotation.

##### Whole genome resequencing of a cross between C1-5 (Female Vitis sylvestris) and Martigny_2 (Male Vitis sylvestris)

These two accessions were crossed in the Inra Colmar ampelographic collection. The accession C1-5 female *Vitis sylvestris* (plant accession number at Inra Colmar 8500.Col.C1-5) originated from the geographic location Sainte-Croix-en- Plaine, Haut-Rhin (68), France, and the male accession Martigny_2 (plant accession number at Inra Colmar 8500.Col.1), was an accession originating from the geographic location Martigny, Swiss. The progeny was grown at Inra Colmar greenhouses. The DNA samples of both parents, of five male descendants (44613.Col.5026T, 44613.Col.5028T, 44613.Col.5029T, 44613.Col.5033T and 44613.Col.5053T) and five female descendants (44613.Col.5035T, 44613.Col.5040T, 44613.Col.5046T, 44613.Col.5050T and 44613.Col.5057T) were sequenced. The quantity and quality of the DNA extracted from leaves (DNeasy plant mini kit, Qiagen) of 12 individuals from the cross (2 parents and 10 offspring) was checked prior to sequencing. Nanodrop analysis indicated that concentrations were always above 110 ng.µl^-1^, and that more than 5 µg DNA was available for all samples. Quality (fragment size) was checked using capillary electrophoresis (Fragment Analyzer) and was satisfactory. Based on quality check, we adapted sonication (using Covaris E220) in order to get mostly fragments of 250 bp. Twelve Illumina libraries were constructed and were pooled for two lanes of sequencing on an Illumina Hiseq 4000 machine in 2×100bp paired-end mode. This yielded between 85 and 149 millions reads per individual (Supplementary Table 2), which roughly corresponded to 17X to 30X coverage of re-seq data per individual.

#### Genome assembly, cleaning, polishing, anchoring and ordering

Shortly, the Sylvestris C1-2 *de novo* genome assembly was performed with Falcon-integrate (Falcon + Falcon-unzip), in order to obtain a diploid assembly (the two haplotypes). With Falcon, PacBio reads were self-corrected, assembly was performed, haplotypes were generated, assembly was polished with Arrow and finally we obtained a phased diploid assembly of Sylvestris C1-2.

##### Genome assembly and phasing

The FALCON-integrate 1.8.4 tool used is available on github: https://github.com/PacificBiosciences/FALCON-integrate/tree/1.8.4. The FALCON parameters used for the *Vitis sylvestris* genome assembly are taken from the genome assembly of *Vitis vinifera* cv Cabernet Sauvignon paper^7^ (pa_HPCdaligner_option = -v -dal128 -e0.75 - M60 -l2500 -k18 -h1250 -s100 -w8; ovlp_HPCdaligner_option = -v -dal128 -M60 -e.96 - l1500 -s100 -k24 -h1250; pa_DBsplit_option = -a -x500 -s200; ovlp_DBsplit_option = -s200; falcon_sense_option = --output_multi --min_idt 0.70 --min_cov 4 --max_n_read 400 --n_core 8; falcon_sense_skip_contained = False; overlap_filtering_setting = --max_diff 120 -- max_cov 120 --min_cov 4 --bestn 10 --n_core 8). The phasing and haplotypes creation was performed with Falcon-unzip, with default parameters. The assembly was polished with FALCON and PacBio reads using Arrow (available in Falcon-integrate).

##### Assembly polishing and finishing

After the FALCON run, we performed additional polishing with Illumina reads. First, illumina reads were aligned with bwa^32^ mem and -M option on FALCON’s genome assembly. Then, alignments were filtered with samtools to keep only primary alignments and concordant pairs. Finally, alignments were filtered with bamtools to keep alignments with an edit distance <= 5. These filtered aligned reads are used to polish the assembly with the pacbio-util (version 0.2) from pacbio-utilities tool (https://github.com/douglasgscofield/PacBio-utilities). Then, the same alignment was performed on the genome polished with pacbio-utils and this genome was polished with Illumina reads and PILON (v1.22 - https://github.com/broadinstitute/pilon). A few haplotigs may have remained in the primary contigs file. A tool is available, purge_haplotigs, to find these false primary contigs in order to move them to the haplotigs file. We used this tool to correct this and to finish our genome asssembly (v1.0.4 - commit 6414f68 - https://bitbucket.org/mroachawri/purge_haplotigs/src/master/).

##### Quality control of the assembly

Assembly statistics such as number of contigs, N50 and L50 were calculated with a home-made script. Genome assembly completeness was assessed with BUSCO^33^ with the genome mode, the embryophyta_odb9 lineage and the *Arabidopsis* species options (version 2.0 - https://gitlab.com/ezlab/busco). Nucmer tool (from MUMmer tool: https://sourceforge.net/projects/mummer/ - nucmer (version 3.1) parameters: -maxmatch –l 100 -c 500) and the grapevine reference genome^8^ (PN40024, version 12X.2 - https://urgi.versailles.inra.fr/Species/Vitis/Data-Sequences/Genome-sequences) were used to align the Sylvestris assembly to the PN40024 reference assembly in order to see completeness of this Sylvestris assembly. Nucmer and grapevine reference genome were also used to anchor and order Sylvestris contigs into chromosomes (with additional parameters –r –q), as it was done in the Chardonnay genome assembly paper^34^.

#### BAC generation and sequencing

##### Generation of BAC library of a Vitis sylvestris male sequence

To obtain the DNA sequence of the Y allele in the sex region, we proceeded as follows. We chose a *Vitis vinifera* ssp. *sylvestris* male from a wild population spontaneously growing on a hill forest near Montpellier (France), on the Northern slope of the Pic Saint Loup mountain. Its male phenotype was confirmed over five years of observation of the flowers, both on the forest plant and on 5 of its clones planted in the INRA Vassal Grape Collection in Marseillan, France (introduction name: Lambrusque PSL10; introduction number: 8500Mtp107). High Molecular Weight DNA was isolated from 40 g of PSL10 young leaves following the cell nuclei extraction method described in Peterson *et al.*^35^ (2000) with slight modifications^36^. The long DNA fragments were partially digested with EcoRI restriction enzyme, and fragments from 100 to 250 kb were selected. Sized and eluted DNA was then ligated into a pAGIBAC- EcoRI cloning vector and cloned into DH10B T1R *Escherichia coli* strain (Invitrogen). The resulting BAC clones were plated on a solid selective medium and organised in barcoded microplates using a robotic workstation QPix2 XT (Molecular Devices). The BAC library was named Vsy-B-Lamb. It consists of 27,648 clones of 113kb size in average, representing a 6.4x genome equivalent coverage. The library was replicated for security reason and the two copies were stored in separate freezers at −80°C (resource available at https://cnrgv.toulouse.inra.fr/fr).

##### BAC library screening

The bacterial clones were deposited on a macroarray nylon membrane (22×22 cm), following a 6×6 grid pattern. Three copies of this gridded macroarray were created. To select BAC clones carrying the DNA fragments from the sex locus, we used specifically designed radio-labelled probes. These probes were defined using the sequences published in Picq et al (2014)^5^: VSVV006, VSVV007, VSVV009, VSVV010, VSVV011 (GeneID GSVIVT01001275001, GSVIVT01001277001, GSVIVT01001286001, GSVIVT00007310001, GSVIVT00007312001 respectively). The hybridisation of 3 separate pools of probes allowed to spot around 200 putatively positive BAC clones in total; the 30 clones showing the most intense spots were individually tested via Real-Time PCR using the same sequences used for probe design and 9 clones were validated. The same clustering of the 9 positive clones into two groups of alleles was obtained using two approaches: the first assignation based on their melting temperature curves similarities; and the second based on Sanger sequencing of internal and BAC-end sequences. These two groups correspond to the two alternate alleles expected in a “XY-like” sex region. Internal and BAC-ends sequences were also used to map the BACs on each other, for each allele, with the objective to sequence the minimal number of clones with an optimised overlapping to cover the whole region (minimum tilling paths composed of 3 and 4 clones respectively).

##### BAC sequencing and assembly of X and Y haplotypes

The sequences of the 7 sex-region BACs of *V. sylvestris* were obtained via a PacBio RSII sequencer (P6C4 chemistry). Sequencing was done in a pool of 20 individually tagged BAC clones. We used 2 µg of each individual BAC clone DNA to prepare the PacBio SMRT® 10kb library. GeT-PlaGe Genomic Platform (INRA-Toulouse, France) handled the sample loading on the RSII device and the data retrieval. We performed the detection and removal of residual *E. coli* sequences on raw reads. After a second cleaning step consisting in detecting and removing the vector sequences, individually-tagged BAC sequences were assembled with the HGAP workflow (https://github.com/PacificBiosciences/Bioinformatics-Training/wiki/HGAP).

For each of the two groups of BACs identified as above, the BAC sequences were joined using their overlapping, so to form a long “haplotig”. One haplotig was then assigned to the “male haplotype” and the other to the “female haplotype” using the sex-discriminating polymorphisms described by Picq et al^5^ (2014), namely 10 SNPs for VSVV006, 7 SNPs for VSVV007 and 6 SNPs for VSVV009. These SNPs were found to be 100% associated with sex, in a worldwide collection of 22 males, 23 hermaphrodites and 91 females (Picq et al^5^ 2014), and we confirmed that the two haplotigs had either all male SNPs or all female SNPs.

#### Genome and BAC annotation

##### Genome annotation

RNA Illumina paired-end reads obtained from grape berries of Sylvestris C1-2 (see above) were used to assemble a transcriptome in order to annotate the genome. Prinseq-lite^37^, Ribopicker^38^ and STAR^39^ were used to clean the data, and the assembly was made using Trinity^40^. We assembled 398,189 putative transcripts (422,652,106 bp), with a N50 of 2,217 bp, which roughly corresponds to the average size of Pinot noir transcripts (2,056 bp). TransDecoder^40^ detected 187,734 CDS with an average size of 832 bp (of which 60,985 CDS are larger than 832 bp). A BUSCO analysis detected 1,237 genes, of which 1,166 (94%) are complete. EuGene-EP^41^ (Eukaryote Pipeline - v1.4 - http://eugene.toulouse.inra.fr/) was used to perform genes annotation on the Sylvestris assembly. The protein databases used for the genes annotation were the TAIR10, swissprot and uniprot plants databases. The transcriptomes used for the genes annotation were our Sylvestris transcriptome, 812 manual annotated genes, and transcript sequences of *Vitis vinifera* from NCBI - 2017-11-08. The configuration file with all the parameters used is available as supplementary data (egnep.Vvi.for_Sylvestris.cfg). Primary contigs and haplotigs were annotated separately, with the same parameters. Finally, we obtained two genes annotations, one for primary contigs and one for haplotigs, with different files: genes annotation in a gff3 file, gene/mrna/cds/ncrna/protein sequences in separate fasta files and some statistics per gene.

##### BAC annotation

The BAC sequences (*V. sylvestris* Male haplotype P2, *V. sylvestris* Male haplotype P1, *V. sylvestris* Female haplotype, *V. vinifera* Cabernet Sauvignon Hermaphrodite genome sequences and the annotation with EuGene^41^ (Eukaryote Pipeline - v1.4 - http://eugene.toulouse.inra.fr/) was started, with same parameters and reference files than for the genome annotation. The protein databases used for the genes annotation were the TAIR10, swissprot and uniprot plants databases. The transcriptomes used for the genes annotation were our Sylvestris transcriptome, 812 manual annotated genes, and transcript sequences of *Vitis vinifera* from NCBI - 2017-11-08. The configuration file with all the parameters used is available as supplementary data (egnep.Vvi.for_Sylvestris.cfg). Then the annotations specific to BACs were extracted and used as final BACs annotation. We did not run Eugene on the 5 BAC sequences alone because EuGene learns from data and we did not want to introduce a bias due to the low number of BAC sequences.

#### Development of SEX-DETector++

We developed a new version of the SEX-DETector software, that implements a probabilistic method to study SNPs segregation in a family, taking into account genotyping errors. SEX-DETector++ is coded in C++ and uses additional algorithmic optimisation to reduce the running time and memory usage by about two orders of magnitude compared to the original code. SEX-DETector++ thus allows convenient usage on large, genome-wide genotyping datasets for which the original code would require prohibitively large running times and memory. The underlying model is the same as in the original code, but new functionalities were added to deal with genomic data as an input of the method (in the vcf format). The code is publicly available at https://gitlab.in2p3.fr/sex-det-family/sex-detector-plusplus, where technical and installation details can also be found.

#### Characterization of the sex-linked region in the *V. sylvestris* genome

##### SNP-tolerant mapping and SNP discovery from whole-genome resequencing data

Divergence between X and Y alleles can prevent SNP discovery. To avoid this issue, we carried out an iterative SNP-tolerant mapping similarly to the procedure described in Prentout *et al.*^11^. Raw reads of twelve individuals of the same family (2 parents and 5 descendants of each sex) were first mapped against the *V. sylvestris* genome with gsnap^42^ v2018-07-04 (-m 0.1) in standard mode. At each iteration step, SNP calling was performed: variants were called with samtools mpileup v1.3.1^43^ followed with Varscan v2.4.3^44^ mpileup2snp (Min coverage 8, Min reads2 2, Min var freq 0.2, Min avg qual 15, P-value thresh 0.01). Additional filters were applied: minimum frequency of variants reads between 0.25 and 0.75 for heterozygote genotypes in individual samples; maximum coverage of twice mode of the gaussian distribution for each sample, only bi-allelic SNPs, minor frequency variant higher than 0.05, maximum rate of missing data 0.2. A SNP database was build with gsnap utilities to perform a SNP-tolerant mapping at the next iteration. We carried out two iterations of SNP-tolerant mapping until the rate of discovery of new XY SNPs (see below) became low. Finally, a 4^th^ step of SNP-tolerant mapping with more stringent parameters (-m 0.05, primary alignments only) was performed in order to reduce the rate of false-positive SNPs.

##### Detection and validation of sex-linked single nucleotide polymorphisms

In order to detect SNPs linked to the sex-determining region, we run SEX-DETector++. This yielded 4,113 sex-linked SNPs (with a posterior probability high than 0.6), 90.5% of them on chromosome 2. As the sex-determining regions is assumed to be small^5^ and the low number of offspring in our cross gives us access to few recombination events, the sex-linked region in our analysis may be larger than the actual sex-determining region (SNP close to the sex locus tend to be linked to sex in a particular cross). To determine the boundaries of the sex-determining region, independent public data were analyzed in order to identify shared XY SNPs between different populations. WG-reseq and RNA-seq data from wild and cultivated grapevines were mapped against the *V. sylvestris* genome with gsnap v2018-07-04, SNPs were called with varscan v2.4.3. We searched for candidate XY-SNPs that were heterozygote in all male samples and homozygote in all female samples of the validation dataset. Validated XY-SNPs spanned 111- kb on chromosome 2 (from 4,810,929 to 4,921,949 bp), which indicated the limits of the sex-determining region. In order to complement SEX-DETector++ analysis, we also performed an empirical one. We searched for SNPs that overlapped the sex-determining region, were homozygote in female individuals of the cross, and heterozygote in male individuals, allowing two missing alleles. This empirical analysis retrieved all 1,406 XY SNPs identified by SEX- DETector++ plus 459 additional ones (+ 32.6%). This final dataset of 1,865 XY SNPs was used for downstream analyses.

##### Determination of the synonymous divergence between X and Y alleles in the sex-linked region

The dataset of XY SNPs was used to build X and Y allelic pseudosequences in coding regions, using a custom python script to substitute reference positions by X or Y SNPs respectively in the genome in respect with strand, and to extract and concatenate coding DNA sequences for each gene of the sex locus. The yn00 program of the PAML suite^45^ was used to estimate synonymous divergence (dS) with standard error estimation.

##### Structural characterization of the sex-linked region in the V. sylvestris genome

Structural gene annotations in the sex-linked regions were extracted from Eugene (See annotation Section). Predicted proteins were mapped against the ncbi nr database with blastp and functional annotations were manually analyzed. The best hit in *Arabidopsis thaliana* was used to name the genes. Transposable elements were detected with Red^46^.

#### Integrative search for sex-determining genes in the sex locus

##### Comparison of gene and TE content in X, Y and H haplotypes

In addition to the female whole genome of *V. sylvestris*, we obtained assemblies for BACs of X and Y haplotypes of *V. sylvestris* and H and F haplotypes of Cabernet-Sauvignon (cf. BAC sequencing and assembly Section), H designating modified Y haplotypes found in hermaphrodites. Blastn (ncbi-blast v2.2.3) were carried out between pairs of haplotypes (X-Y, X-H, Y-H, X from whole-genome and X from BACs). Whole-haplotype comparisons were visualized with circos to identify presence/absence patterns. Genic sequences were extracted in all haplotypes using Eugene annotations and blastn, and aligned with mafft v7 online service^47^ to identify structural differences and mutations. The sequences of the putative sex-determining candidates were blasted against assemblies of other cultivated grapevine genomes (Chardonnay, Cabernet-Sauvignon and Zinfandel) to assess their presence or absence.

##### Total and allele-specific expression of sex-linked genes

We retrieved raw reads from 23 public libraries of female, male and hermaphrodite bud flowers samples from *V. sylvestris* at four developmental stages^13^. Reads were mapped on the *V. sylvestris* genome with gsnap v2018-07-04 (-m 0.1), with a mode tolerant to XY SNPs. Read counts were obtained with htseq-count v0.10.0^48^ and normalized by computing the RPKM (reads per count per millions of reads mapped). To specifically measure the expression of X and Y alleles, we performed SNP calling (gsnap –m 0.05, varscan max missing data 0.4 and minor allele frequency 0.05), and extracted the positions corresponding to XY SNPs located in CDS from the vcf file. For each gene and library, we read the number of reads mapping on reference and variant alleles (corresponding to X and Y alleles respectively) with bedtools interesect^49^. Reads numbers were summed by gene and normalized by the total number of reads in the variant file for the library and the length of transcripts.

##### Organ-specific expression of sex-linked genes in V. vinifera

We retrieved and mapped transcriptomic data of *V. sylvestris* and *V. vinifera* in several organs and conditions against the genome of *V. sylvestris* (berries, developing seeds, leaves under normal, drought and pathogenic conditions, stem, early and mature flower buds, Supplementary Table 12). We measured the gene expression levels in the different conditions, and computed an index of organ-specificity (Tau^50^) on normalized expression levels (log(read count per kilobase per millions of mapped reads)). The Tau specificity index ranges between 0 and 1 and typically display two modes near 0 and 1 indicating ubiquitous and organ-specific genes respectively.

## Supplementary Figures

**Supplementary Figure 1:**
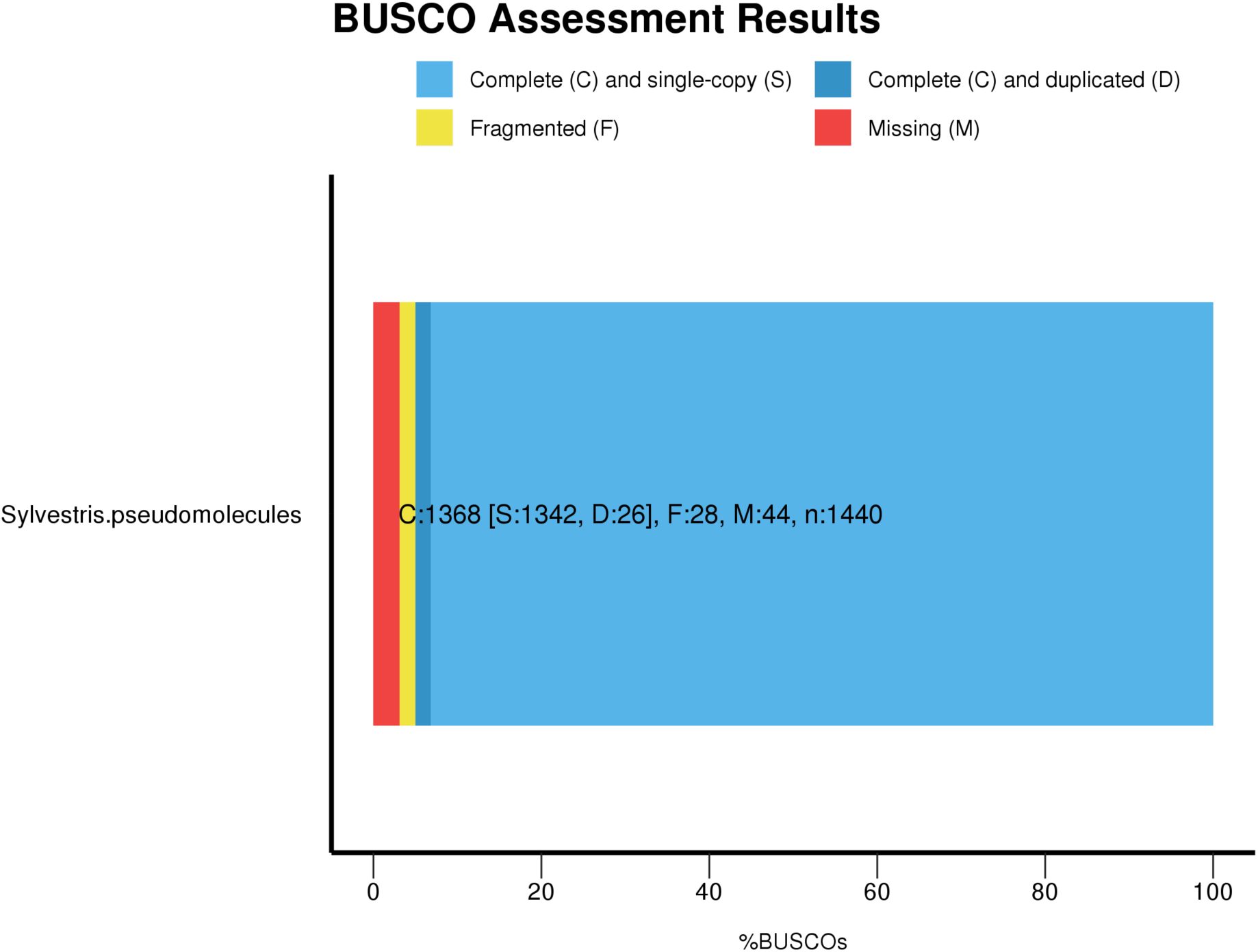
BUSCO results on *V. sylvestris* pseudomolecules. Out of 1440 genes in the BUSCO dataset, 95% are complete.

**Supplementary Figure 2:**
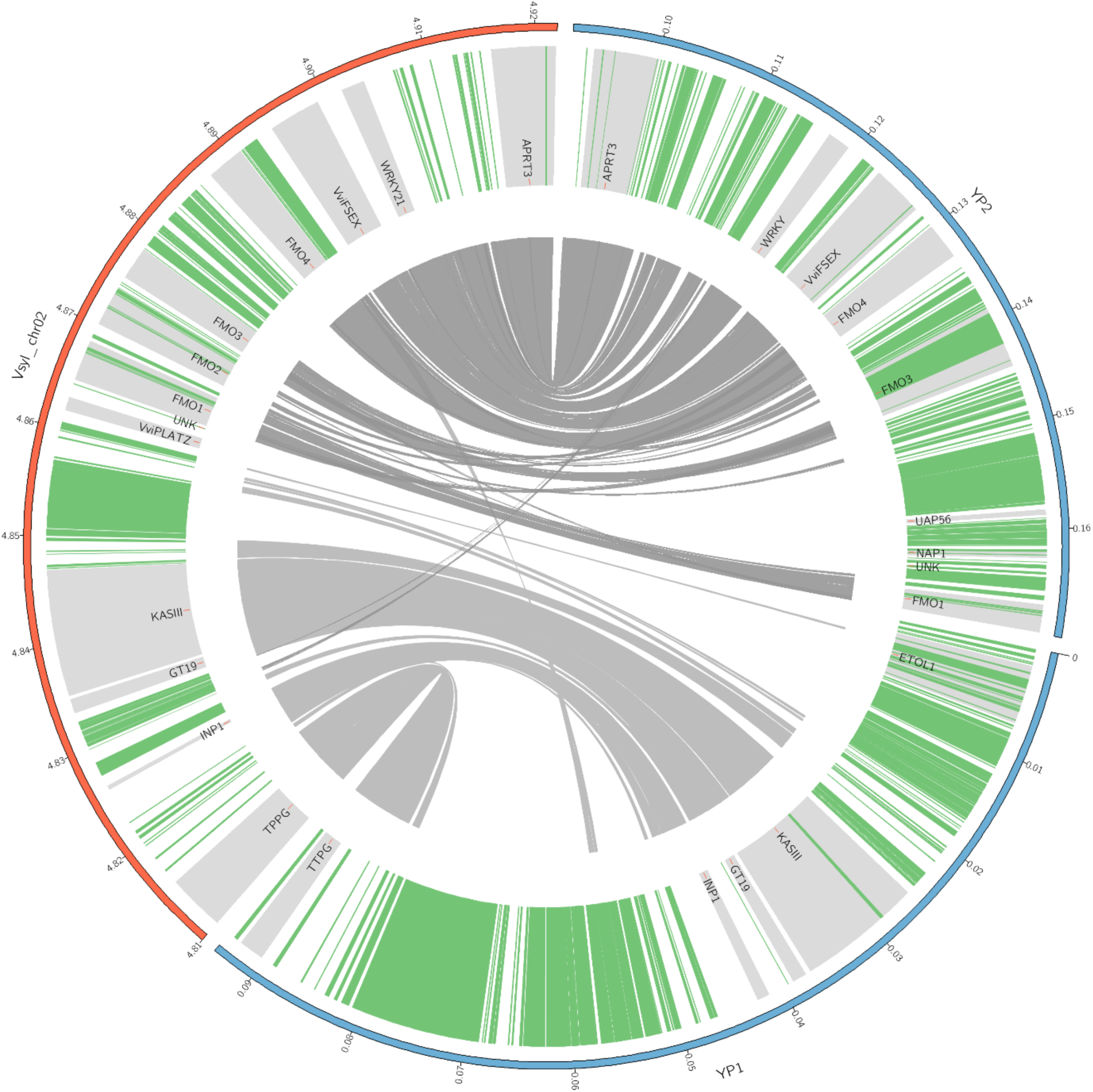
Structural comparison of X and Y haplotypes of the sex locus of *V. sylvestris*. Outer to inner track: circular representation of pseudomolecules; limits of genes (obtained from Eugene and verified with blastn) in grey and repeats in green; synteny relationships (blastn hits with an e-value lower than 0.01. YP1 and YP2 are two BAC contigs covering the Y locus, with a gap of an estimated size of 13kb, in which presence of the *PLATZy* allele has been confirmed by PCR (see Supplementary Text). Coordinates are indicated in Mbp.

**Supplementary Figure 3:**
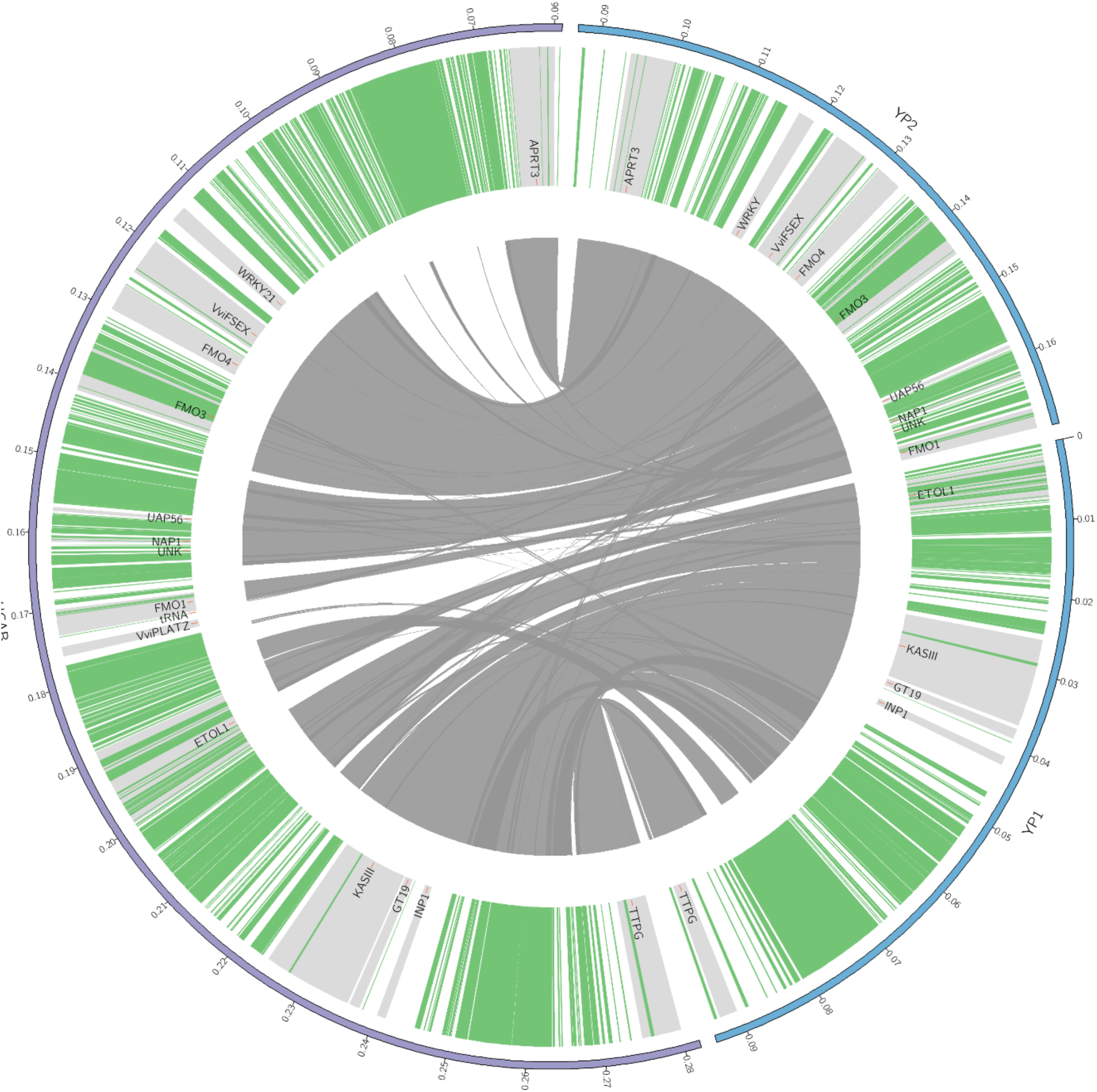
Structural comparison of a Y haplotype of *V. sylvestris* and a H haplotype of *V. vinifera* cv. Cabernet Sauvignon, H referring to the modified Y haplotype found in hermaphrodites. Outer to inner track: circular representation of pseudomolecules; limits of genes (obtained from Eugene and verified with blastn) in grey and repeats in green; synteny relationships (blastn hits with e-values lower than 0.001). YP1 and YP2 are two BAC contigs covering the Y locus, with a gap of an estimated size of 13kb, in which presence of the *PLATZy* allele has been confirmed by PCR (see Supplementary Text). The purple ideogram represents the H haplotype. A large insertion between *WRKY* and *APRT3* is present in the H haplotype of Cabernet Sauvignon, but is not shared by all H haplotypes (*e.g.* only of the two H haplotypes of Chardonnay, not shown). Coordinates are indicated in Mbp.

**Supplementary Figure 4:**
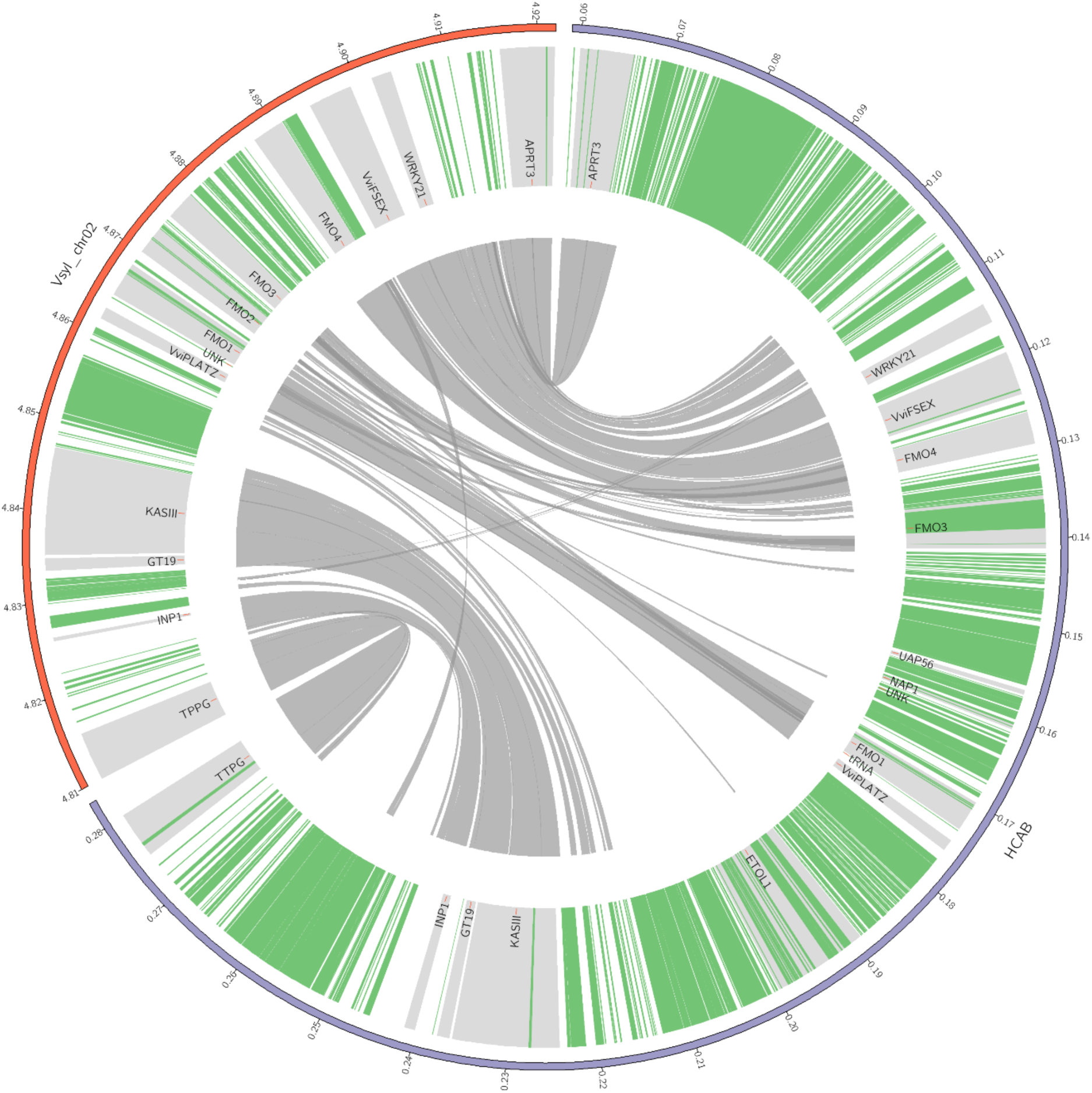
Structural comparison of X haplotype of *V. sylvestris* and a H haplotype of *V. vinifera* cv. Cabernet Sauvignon. Outer to inner track: circular representation of pseudomolecules; Outer to inner track: circular representation of pseudomolecules; limits of genes (obtained from Eugene and verified with blastn) in grey and repeats in green; synteny relationships; synteny relationships (blastn hits with e-values lower than 0.001). Coordinates are indicated in Mbp.

**Supplementary Figure 5:**
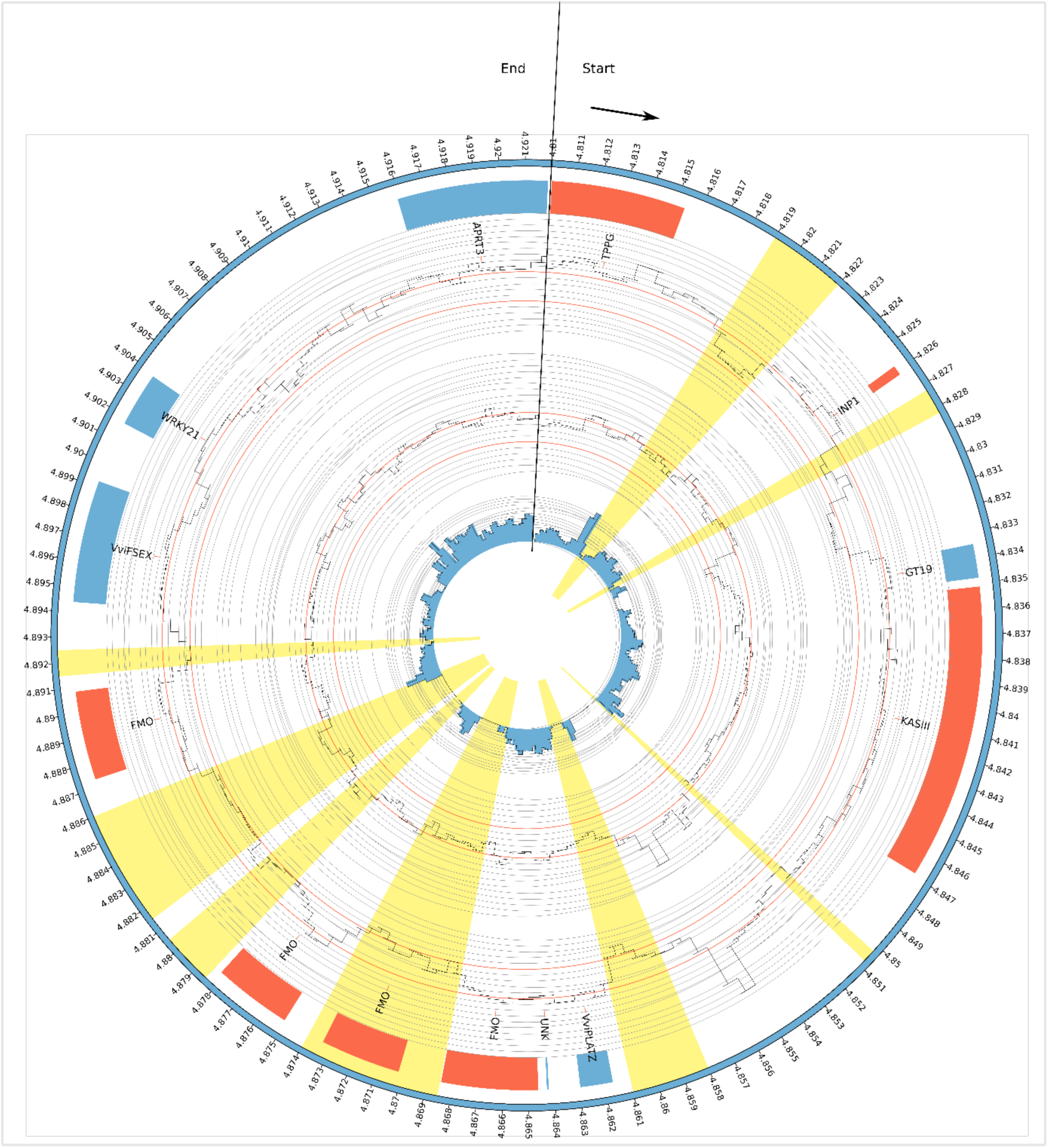
Density of XY-SNPs and location of X-hemizygote regions in a cross in *V. sylvestris*. Track, form outter to inner track: genomic coordinates on chromosome 2 (from 4.810 to 4.922 Mb), location and name of genes (blue = direct strand, red = reverse strand), Depth of mapping coverage in a male individual (RCDN9), from 0 to 50, Depth of mapping coverage in a female individual (RCDN16), number of XY SNPs (from 0 to 64). All values were computed by 1-kb overlapping windows. The number of XY SNP was not normalized in order to reflect the proportion of informative sites in each window. Yellow highlights represent the approximate limites of X-hemizygote regions inferred from a two-fold reduction of mapping coverage in males.

**Supplementary Figure 6:**
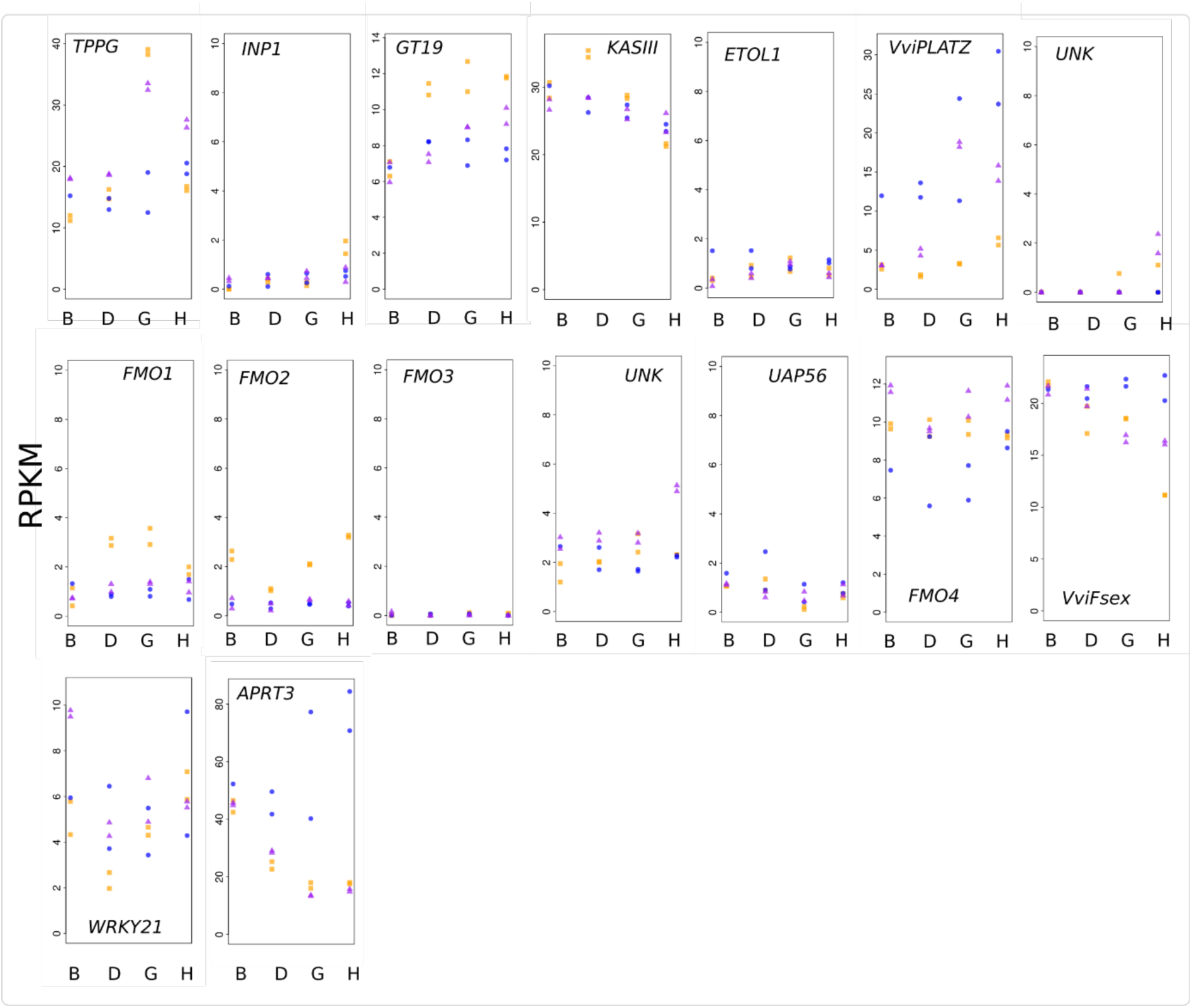
Total normalized gene expression in males, females and hermaphrodites of *V. sylvestris* for genes in the sex locus during flower bud development. Letters B to H reffer to successive developmental stages. Expression in females, males and hermaphrodites are represented by orange, purple and blue colors, respectively.

**Supplementary Figure 7:**
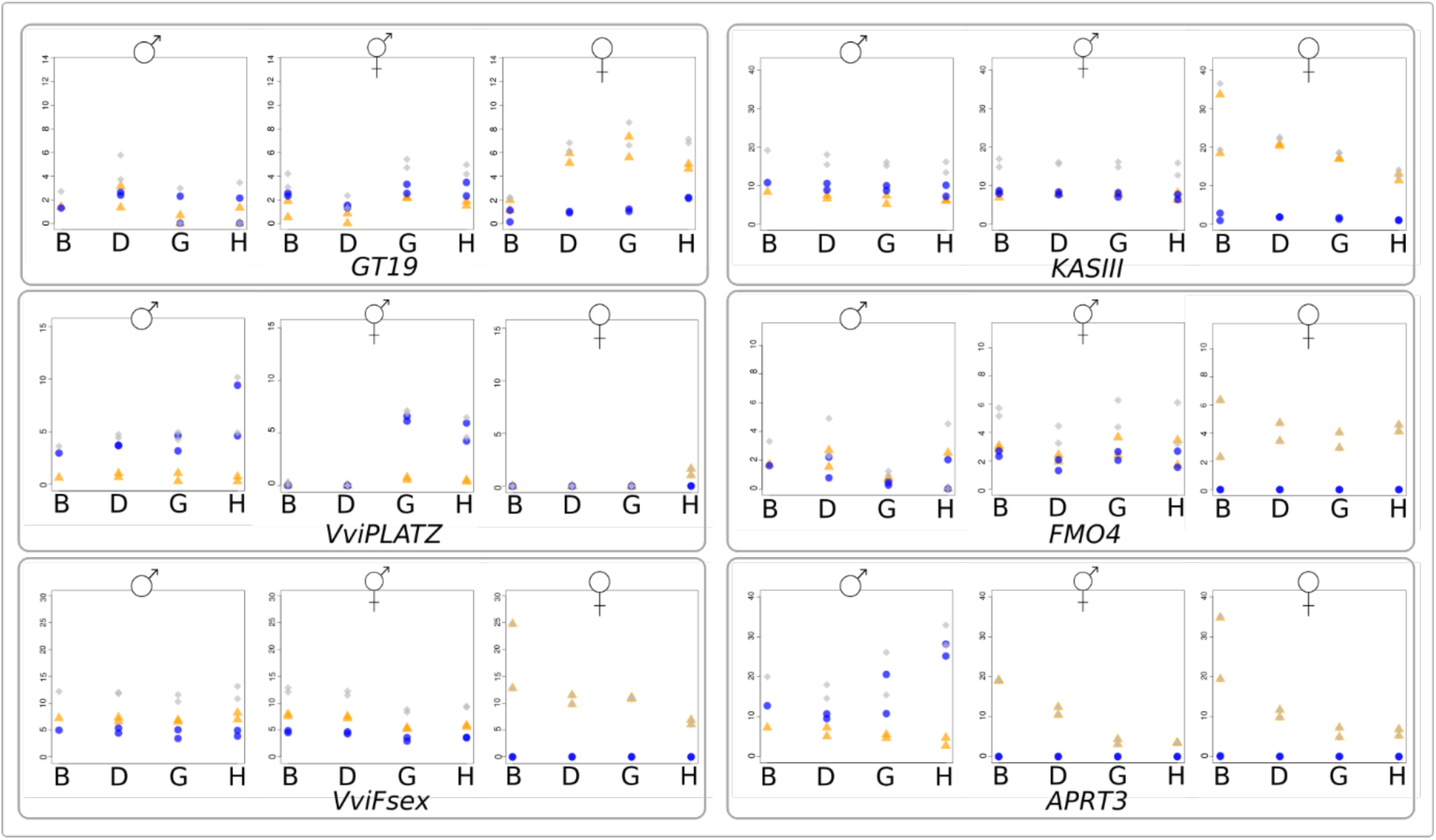
Allele-specific expression of X and Y alleles of XY genes pairs of *Vitis sylvestris* during the development of flower buds. The X and Y allelic expression from B to H-stages flower buds is shown for females, males and hermaphrodites. Orange triangles, blue dots and grey losanges represent X, Y and total allelic expression respectively. Allele-specific expression was computed only for genes with sufficient mapping coverage on at least 5 XY SNPs. The y scale is different for each gene but shared for a given gene between females, males and hermaphrodites. Two replicates are represented for each condition, except for the male B stage for which there was no replicate.

**Supplementary Figure 8:**
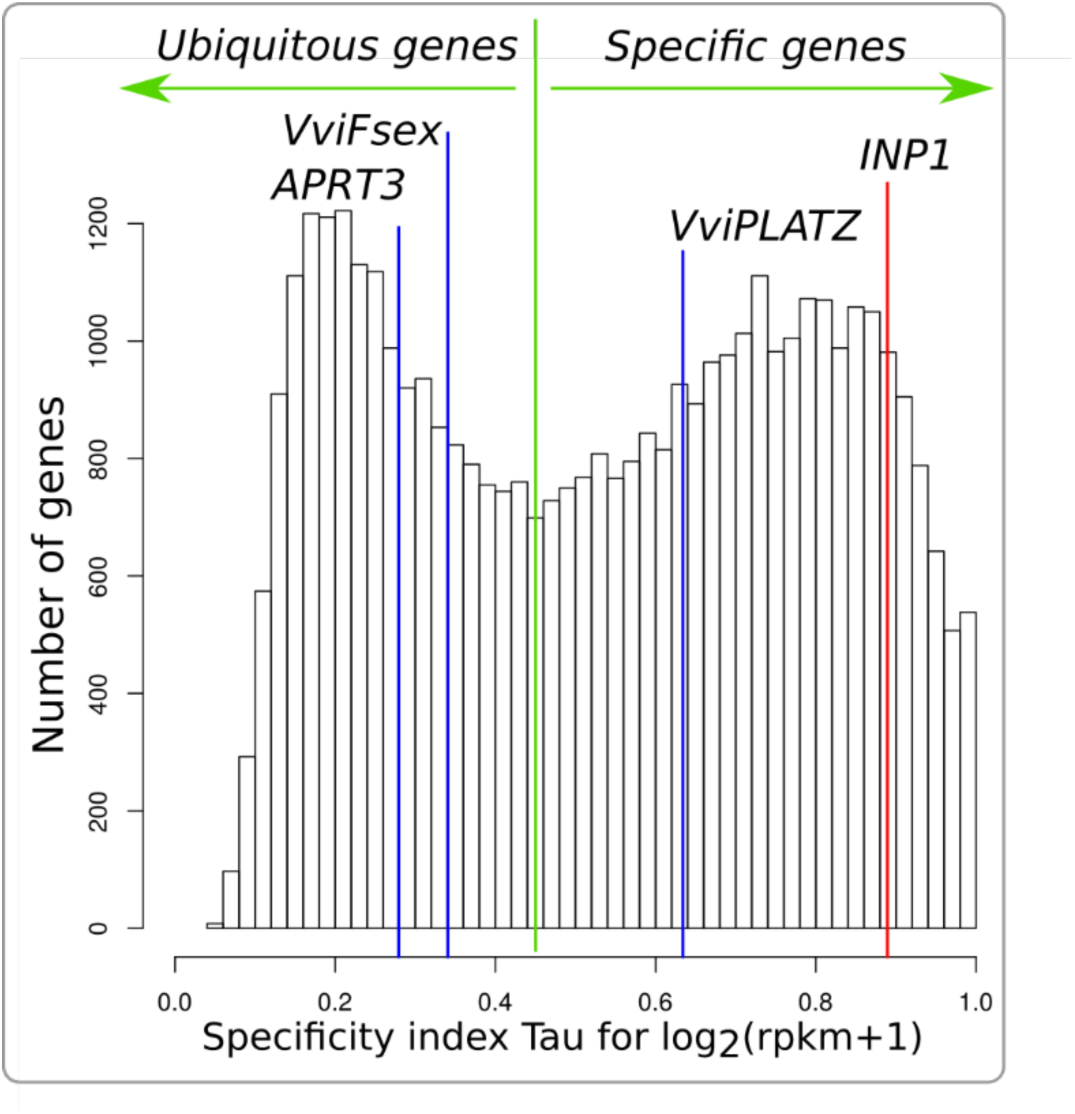
Distribution of organ-specificity expression in *Vitis sylvestris* genes. The value of the specificity index Tau for four genes of the sex locus in indicated by vertical lines.

## Supplementary Tables

**Supplementary Table 1:** Assembly and anchoring statistics of the *V. sylvestris* genome.

|  | Primary contigs | Anchored primary contigs | Haplotigs |
| --- | --- | --- | --- |
| Number of contigs | 591 | 476 | 3,781 |
| Mean length (b) | 792,724 | 947,992 | 83,879 |
| Median length (b) | 421,056 | 584,494.5 | 36,553 |
| Max length (b) | 6,865,695 | 6,865,695 | 2,541,772 |
| L50 | 82 | 77 | 483 |
| N50 (b) | 1,711,677 | 1,773,898 | 173,606 |
| Assembly size (b) | 468,500,071 | 451,244,463 | 317,149,633 |

**Supplementary Table 2:** Summary statistics of resequencing data of a cross between two *V. sylvestris* parents.

| Sample ID | Genotype | Type | Sex | Concentration ADN ng/μl | Number of Reads | Number of Pairs |
| --- | --- | --- | --- | --- | --- | --- |
| RCDN8 | C1-5 | mother | Female | 158 | 91,689,312 | 45,844,656 |
| RCDN9 | Martigny 2 | father | Male | 113 | 140,875,256 | 70,437,628 |
| RCDN10 | 5026T | offspring | Male | 150 | 149,542,818 | 74,771,409 |
| RCDN11 | 5028T | offspring | Male | 145 | 128,590,900 | 64,295,450 |
| RCDN12 | 5029T | offspring | Male | 145 | 85,976,170 | 42,988,085 |
| RCDN13 | 5033T | offspring | Male | 145 | 130,071,908 | 65,035,954 |
| RCDN14 | 5053T | offspring | Male | 155 | 123,960,284 | 61,980,142 |
| RCDN15 | 5035T | offspring | female | 110 | 109,564,824 | 54,782,412 |
| RCDN16 | 5040T | offspring | female | 110 | 132,609,212 | 66,304,606 |
| RCDN17 | 5046T | offspring | female | 130 | 111,932,706 | 55,966,353 |
| RCDN18 | 5050T | offspring | female | 110 | 106,023,554 | 53,011,777 |
| RCDN19 | 5057T | offspring | female | 120 | 125,195,484 | 62,597,742 |

**Supplementary Table 3:** Mode of the gaussian distribution of mapping coverage for each sample for the dataset of whole genome resequencing of a cross in *V. sylvestris*. The values are identical for each iteration of mapping and therefore indicated only once. The maximum coverage allowed corresponds to the maximum coverage that was allowed when filtering SNP, in order to remove repeated positions.

| Sample ID | Mode of gaussian distribution | Maximum coverage allowed |
| --- | --- | --- |
| RCDN8 | 16 | 32 |
| RCDN9 | 25 | 50 |
| RCDN10 | 26 | 52 |
| RCDN11 | 22 | 44 |
| RCDN12 | 14 | 28 |
| RCDN13 | 22 | 44 |
| RCDN14 | 21 | 42 |
| RCDN15 | 19 | 38 |
| RCDN16 | 23 | 46 |
| RCDN17 | 19 | 38 |
| RCDN18 | 18 | 36 |
| RCDN19 | 22 | 44 |

**Supplementary Table 4:** Summary statistics of iterative SNP-tolerant mapping for the resequencing dataset of a cross in *V. sylvestris*.

| Sample ID | Sex | Iteration 1 |  |  | Iteration 2 |  |  |
| --- | --- | --- | --- | --- | --- | --- | --- |
|  |  | Number of mapped reads | fraction of mapped reads | fraction of paired reads | Number of mapped reads | fraction of mapped reads | fraction of paired reads |
| RCDN8 | F | 89,951,765 | 0.981 | 0.969 | 90,089,021 | 0.983 | 0.971 |
| RCDN9 | M | 138,338,630 | 0.982 | 0.971 | 138,544,634 | 0.983 | 0.973 |
| RCDN10 | M | 146,671,291 | 0.981 | 0.968 | 146,904,980 | 0.982 | 0.971 |
| RCDN11 | M | 125,913,524 | 0.979 | 0.966 | 126,122,927 | 0.981 | 0.969 |
| RCDN12 | M | 84,214,280 | 0.98 | 0.967 | 84,352,679 | 0.981 | 0.969 |
| RCDN13 | M | 127,437,731 | 0.98 | 0.967 | 127,645,009 | 0.981 | 0.969 |
| RCDN14 | M | 121,369,732 | 0.979 | 0.966 | 121,578,081 | 0.981 | 0.968 |
| RCDN15 | F | 107,360,998 | 0.98 | 0.967 | 107,538,009 | 0.982 | 0.97 |
| RCDN16 | F | 130,211,584 | 0.982 | 0.971 | 130,408,313 | 0.983 | 0.973 |
| RCDN17 | F | 109,719,948 | 0.98 | 0.968 | 109,900,170 | 0.982 | 0.97 |
| RCDN18 | F | 104,161,079 | 0.982 | 0.971 | 104,315,801 | 0.984 | 0.973 |
| RCDN19 | F | 122,729,069 | 0.98 | 0.967 | 122,927,332 | 0.982 | 0.97 |

| Sample ID | Iteration 3 |  |  | Iteration 4 |  |  |
| --- | --- | --- | --- | --- | --- | --- |
|  | Number of mapped reads | fraction of mapped reads | fraction of paired reads | Number of mapped reads | fraction of mapped reads | fraction of paired reads |
| RCDN8 | 90,095,720 | 0.983 | 0.972 | 90,029,816 | 0.982 | 0.97 |
| RCDN9 | 138,552,746 | 0.984 | 0.973 | 138,452,568 | 0.983 | 0.972 |
| RCDN10 | 146,916,364 | 0.982 | 0.971 | 146,801,740 | 0.982 | 0.969 |
| RCDN11 | 126,131,254 | 0.981 | 0.969 | 126,036,403 | 0.98 | 0.968 |
| RCDN12 | 84,358,053 | 0.981 | 0.969 | 84,294,600 | 0.98 | 0.968 |
| RCDN13 | 127,653,125 | 0.981 | 0.969 | 127,558,702 | 0.981 | 0.968 |
| RCDN14 | 121,585,814 | 0.981 | 0.968 | 121,490,591 | 0.98 | 0.967 |
| RCDN15 | 107,545,495 | 0.982 | 0.97 | 107,458,696 | 0.981 | 0.968 |
| RCDN16 | 130,416,024 | 0.983 | 0.973 | 130,319,644 | 0.983 | 0.972 |
| RCDN17 | 109,907,858 | 0.982 | 0.97 | 109,819,091 | 0.981 | 0.969 |
| RCDN18 | 104,322,423 | 0.984 | 0.973 | 104,244,590 | 0.983 | 0.972 |
| RCDN19 | 122,935,210 | 0.982 | 0.97 | 122,840,655 | 0.981 | 0.969 |

**Supplementary Table 5:** Summary statistics of missingness per sample for each iteration of SNP- calling in the dataset of whole-genome resequencing of a cross in *V. sylvestris*. Statistics were obtained with vcftools version 0.1.15. N_MISS is the number of missing sites, F_MISS is the frequency of missing sites.

| Sample ID | Iteration 1 |  | Iteration 2 |  | Iteration 3 |  | Iteration 4 |  |
| --- | --- | --- | --- | --- | --- | --- | --- | --- |
|  | N_MISS | F_MISS | N_MISS | F_MISS | N_MISS | F_MISS | N_MISS | F_MISS |
| RCDN8 | 280,469 | 0.0685 | 257,255 | 0.0576 | 260,910 | 0.0557 | 259,983 | 0.0556 |
| RCDN9 | 80,415 | 0.0196 | 78,141 | 0.0175 | 82,728 | 0.0177 | 82,577 | 0.0177 |
| RCDN10 | 34,051 | 0.00831 | 45,378 | 0.0102 | 55,432 | 0.0118 | 57,368 | 0.0123 |
| RCDN11 | 72,953 | 0.0178 | 79,836 | 0.0179 | 91,745 | 0.0196 | 90,777 | 0.0194 |
| RCDN12 | 418,876 | 0.102 | 385,349 | 0.0863 | 397,437 | 0.0848 | 395,665 | 0.0847 |
| RCDN13 | 59,795 | 0.0146 | 67,214 | 0.0151 | 77,745 | 0.0166 | 78,511 | 0.0168 |
| RCDN14 | 81,071 | 0.0198 | 75,386 | 0.0169 | 80,678 | 0.0172 | 79,763 | 0.0171 |
| RCDN15 | 112,434 | 0.0274 | 103,972 | 0.0233 | 110,945 | 0.0237 | 110,066 | 0.0236 |
| RCDN16 | 51,669 | 0.0126 | 58,007 | 0.0130 | 66,040 | 0.0141 | 66,795 | 0.0143 |
| RCDN17 | 112,887 | 0.0276 | 106,827 | 0.0239 | 114,224 | 0.0244 | 112,998 | 0.0242 |
| RCDN18 | 138,484 | 0.0338 | 133,571 | 0.0299 | 143,660 | 0.0307 | 142,622 | 0.0305 |
| RCDN19 | 54,255 | 0.0132 | 53,412 | 0.0120 | 58,227 | 0.0124 | 57,799 | 0.0124 |

**Supplementary Table 6:**
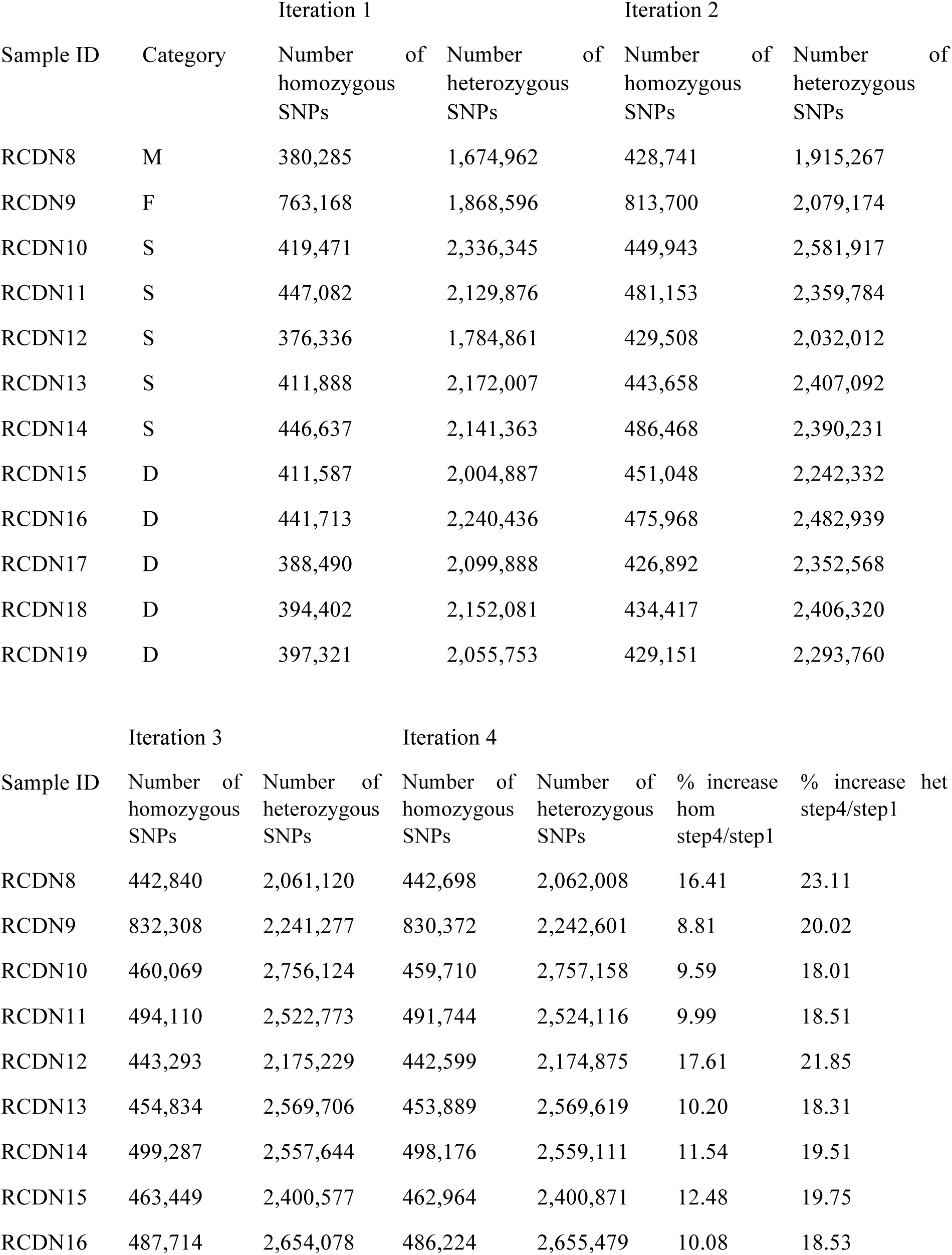
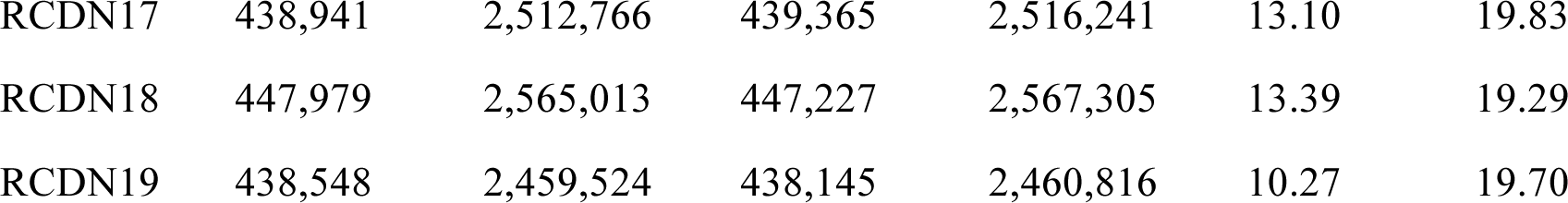
Number of homozygous and heterozygous SNPs called at each iteration in the dataset of whole-genome resequencing of a cross in *V. sylvestris*. “% increase het step4/step1” indicates the percentage of increase in the number heterozygous SNPs detected between the first and fourth step of iterative mapping.

| Sample ID | Category | Iteration 1 |  | Iteration 2 |  |
| --- | --- | --- | --- | --- | --- |
|  |  | Number of homozygous SNPs | Number of heterozygous SNPs | Number of homozygous SNPs | Number of heterozygous SNPs |
| RCDN8 | M | 380,285 | 1,674,962 | 428,741 | 1,915,267 |
| RCDN9 | F | 763,168 | 1,868,596 | 813,700 | 2,079,174 |
| RCDN10 | S | 419,471 | 2,336,345 | 449,943 | 2,581,917 |
| RCDN11 | S | 447,082 | 2,129,876 | 481,153 | 2,359,784 |
| RCDN12 | S | 376,336 | 1,784,861 | 429,508 | 2,032,012 |
| RCDN13 | S | 411,888 | 2,172,007 | 443,658 | 2,407,092 |
| RCDN14 | S | 446,637 | 2,141,363 | 486,468 | 2,390,231 |
| RCDN15 | D | 411,587 | 2,004,887 | 451,048 | 2,242,332 |
| RCDN16 | D | 441,713 | 2,240,436 | 475,968 | 2,482,939 |
| RCDN17 | D | 388,490 | 2,099,888 | 426,892 | 2,352,568 |
| RCDN18 | D | 394,402 | 2,152,081 | 434,417 | 2,406,320 |
| RCDN19 | D | 397,321 | 2,055,753 | 429,151 | 2,293,760 |

| Sample ID | Iteration 3 |  | Iteration 4 |  | % increase hom step4/step1 | % increase het step4/step1 |
| --- | --- | --- | --- | --- | --- | --- |
|  | Number of homozygous SNPs | Number of heterozygous SNPs | Number of homozygous SNPs | Number of heterozygous SNPs |  |  |
| RCDN8 | 442,840 | 2,061,120 | 442,698 | 2,062,008 | 16.41 | 23.11 |
| RCDN9 | 832,308 | 2,241,277 | 830,372 | 2,242,601 | 8.81 | 20.02 |
| RCDN10 | 460,069 | 2,756,124 | 459,710 | 2,757,158 | 9.59 | 18.01 |
| RCDN11 | 494,110 | 2,522,773 | 491,744 | 2,524,116 | 9.99 | 18.51 |
| RCDN12 | 443,293 | 2,175,229 | 442,599 | 2,174,875 | 17.61 | 21.85 |
| RCDN13 | 454,834 | 2,569,706 | 453,889 | 2,569,619 | 10.20 | 18.31 |
| RCDN14 | 499,287 | 2,557,644 | 498,176 | 2,559,111 | 11.54 | 19.51 |
| RCDN15 | 463,449 | 2,400,577 | 462,964 | 2,400,871 | 12.48 | 19.75 |
| RCDN16 | 487,714 | 2,654,078 | 486,224 | 2,655,479 | 10.08 | 18.53 |
| RCDN17 | 438,941 | 2,512,766 | 439,365 | 2,516,241 | 13.10 | 19.83 |
| RCDN18 | 447,979 | 2,565,013 | 447,227 | 2,567,305 | 13.39 | 19.29 |
| RCDN19 | 438,548 | 2,459,524 | 438,145 | 2,460,816 | 10.27 | 19.70 |

**Supplementary Table 7:** Whole-genome resequencing dataset of the ncbi short read archive (SRA) that were mapped to the *Vitis sylvestris* genome. All samples were part of study SRP108271. Sample SRS2238702 had a lower mapping coverage than other samples, resulting in a high rate of missing data.

| SRA run ID | Sample Accession | Domestication status | Organism Name | Origin / cultivar | Sex |
| --- | --- | --- | --- | --- | --- |
| SRR5627780 | SRS2238707 | cultivated | *Vitis vinifera* | Thompson RLK | Hermaphrodite |
| SRR5627781 | SRS2238705 | cultivated | *Vitis vinifera* | Muscat of Alexandria | Hermaphrodite |
| SRR5627782 | SRS2238704 | cultivated | *Vitis vinifera* | Thopson 2A | Hermaphrodite |
| SRR5627783 | SRS2238703 | wild | *Vitis vinifera subsp. sylvestris* | Turkmenistan1 | Malqe |
| SRR5627784 | SRS2238702 | wild | *Vitis vinifera subsp. sylvestris* | Armenia | Female |
| SRR5627785 | SRS2238701 | wild | *Vitis vinifera subsp. sylvestris* | Pakistan1 | Female |
| SRR5627786 | SRS2238700 | wild | *Vitis vinifera subsp. sylvestris* | Turkmenistan2 | Male |
| SRR5627787 | SRS2238699 | wild | *Vitis vinifera subsp. sylvestris* | Azerbaijan1 | Female |
| SRR5627789 | SRS2238698 | wild | *Vitis vinifera subsp. sylvestris* | Georgia | Female |
| SRR5627790 | SRS2238696 | wild | *Vitis vinifera subsp. sylvestris* | Azerbaijan2 | Female |
| SRR5627791 | SRS2238695 | wild | *Vitis vinifera subsp. sylvestris* | Pakistan3 | Female |
| SRR5627792 | SRS2238694 | wild | *Vitis vinifera subsp. sylvestris* | Pakistan2 | Male |
| SRR5627793 | SRS2238693 | cultivated | *Vitis vinifera* | Semillion | Hermaphrodite |
| SRR5627794 | SRS2238692 | cultivated | *Vitis vinifera* | Riesling | Hermaphrodite |
| SRR5627795 | SRS2238691 | cultivated | *Vitis vinifera* | Cabernet Sauvignon | Hermaphrodite |
| SRR5627796 | SRS2238690 | cultivated | *Vitis vinifera* | Primitivo | Hermaphrodite |
| SRR5627797 | SRS2238689 | cultivated | *Vitis vinifera* | Pinot Noir | Hermaphrodite |
| SRR5627798 | SRS2238688 | cultivated | *Vitis vinifera* | Gamay Noir | Hermaphrodite |
| SRR5627799 | SRS2238687 | cultivated | *Vitis vinifera* | Chardonnay | Hermaphrodite |
| SRR5627800 | SRS2238686 | cultivated | *Vitis vinifera* | Aramon | Hermaphrodite |
| SRR5627801 | SRS2238684 | cultivated | *Vitis vinifera* | Zinfandel | Hermaphrodite |
| SRR5627802 | SRS2238685 | cultivated | *Vitis vinifera* | Traminer | Hermaphrodite |

**Supplementary Table 8:** Statistics of mapping and SNP-calling of a public whole-genome resequencing dataset of the ncbi that was mapped to the *Vitis sylvestris* genome.

| SRA run ID | Status | Number of<br>reads | Proportion of<br>mapped reads | Proportion of<br>paired reads |
| --- | --- | --- | --- | --- |
| SRR5627780 | cultivated | 70,284,983 | 0.989 | 0.975 |
| SRR5627781 | cultivated | 76,803,804 | 0.991 | 0.98 |
| SRR5627782 | cultivated | 80,937,324 | 0.992 | 0.981 |
| SRR5627783 | wild | 80,232,477 | 0.986 | 0.974 |
| SRR5627784 | wild | 35,879,048 | 0.954 | 0.92 |
| SRR5627785 | wild | 105,896,803 | 0.991 | 0.978 |
| SRR5627786 | wild | 80,948,588 | 0.99 | 0.979 |
| SRR5627787 | wild | 76,824,067 | 0.985 | 0.97 |
| SRR5627789 | wild | 80,107,055 | 0.989 | 0.977 |
| SRR5627790 | wild | 79,079,895 | 0.965 | 0.938 |
| SRR5627791 | wild | 105,783,610 | 0.991 | 0.979 |
| SRR5627792 | wild | 115,956,970 | 0.991 | 0.976 |
| SRR5627793 | cultivated | 77,221,117 | 0.992 | 0.98 |
| SRR5627794 | cultivated | 77,644,996 | 0.992 | 0.98 |
| SRR5627795 | cultivated | 71,775,105 | 0.992 | 0.98 |
| SRR5627796 | cultivated | 169,763,362 | 0.972 | 0.942 |
| SRR5627797 | cultivated | 90,757,557 | 0.993 | 0.981 |
| SRR5627798 | cultivated | 66,283,833 | 0.992 | 0.983 |
| SRR5627799 | cultivated | 454,053,065 | 0.982 | 0.956 |
| SRR5627800 | cultivated | 96,530,135 | 0.992 | 0.98 |
| SRR5627801 | cultivated | 223,282,216 | 0.992 | 0.978 |
| SRR5627802 | cultivated | 60,767,595 | 0.99 | 0.974 |

**Supplementary Table 9:** Summary of the genotype of 13 cultivars in the sex locus on chromosome 2 inferred from whole-genome resequencing data. Cultivars were genotyped at XY SNPs. H reffers to the modified Y haplotypes in hermaphrodites. Raw SNP data are shown in Figure 1a.

| SRA run ID | Origin / cultivar | Genotype 4.810-4.903 Mb | Genotype 4.903-4.21Mb |
| --- | --- | --- | --- |
| SRR5627780 | Thompson RLK | XH | XH |
| SRR5627781 | Muscat of Alexandria | XH | XH |
| SRR5627782 | Thopson 2A | XH | XH |
| SRR5627793 | Semillion | XH | XH |
| SRR5627794 | Riesling | HH | XH |
| SRR5627795 | Cabernet Sauvignon | XH | XH |
| SRR5627796 | Primitivo | XH | XH |
| SRR5627797 | Pinot Noir | XH | XX |
| SRR5627798 | Gamay Noir | XH | XX |
| SRR5627799 | Chardonnay | HH | XH |
| SRR5627800 | Aramon | XH | XX |
| SRR5627801 | Zinfindel | XH | XH |
| SRR5627802 | Traminer | XH | XX |

**Supplementary Table 10:** Predicted genes in the sex locus of *V. sylvestris*. Coordinates and geneID are indicated for the *V. sylvestris* reference genome, therefore the absolute position of X-deleted genes is not shown.

| Gene ID | Start | End | Strand | Functional annotation | Genotype in hermaphrodites |
| --- | --- | --- | --- | --- | --- |
| 000003Fg0028711 | 4,810,035 | 4,815,364 | - | trehalose phosphate phosphatase | from exon 9, XH or HH |
| 000003Fg0028721 | 4,825,791 | 4,826,183 | - | Protein INAPERTURATE POLLEN1 (INP1) | XH or HH |
| 000003Fg0028731 | 4,833,463 | 4,834,804 | + | xyloglucan galactosyltransferase (GT19) | XH or HH |
| 000003Fg0028741 | 4,835,150 | 4,846,883 | - | 3-oxoacyl-[acyl-carrier-protein] synthase (KASIII) | XH or HH |
| NA |  |  |  | Ethylene-overproducer-like1 / No Pollen Germination 1 (ETOL1) | XH or HH |
| 000003Fg0028751 | 4,861,687 | 4,862,981 | + | transcription of the PLATZ family (VviPLATZ) | XH or HH |
| 000003Fg0028761 | 4,864,219 | 4,864,291 | + | tRNA-gly | XH or HH |
| 000003Fg0028771 | 4,864,636 | 4,868,462 | - | flavin mono-oxygenase (FMO) | XH or HH |
| 000003Fg0028781 | 4,870,175 | 4,873,359 | - | flavin mono-oxygenase (FMO) | XH or HH |
| NA |  |  |  | ATP-dependent DEAD BOX helicase (UAP56) | XH or HH |
| NA |  |  |  | NUCLEOSOME ASSEMBLY PROTEIN 1 (NAP1) | XH or HH |
| NA |  |  |  | Uncharacterized protein (UNK) |  |
| 000003Fg0028791 | 4,875,086 | 4,878,202 | - | flavin mono-oxygenase (FMO) | XH or HH |
| 000003Fg0028801 | 4,887,323 | 4,890,854 | - | flavin mono-oxygenase (FMO) | XH or HH |
| 000003Fg0028811 | 4,894,375 | 4,899,225 | + | Uncharacterized protein (VviFSEX) | XH or HH |
| 000003Fg0028821 | 4,901,701 | 4,903,864 | + | WRKY transcription factor 21 (WRKY21) | XH or HH |
| 000003Fg0028831 | 4,915,923 | 4,921,889 | + | adenine phospho-ribosyltransferase (VviAPRT3) | XH or HH |

**Supplementary Table 11:**
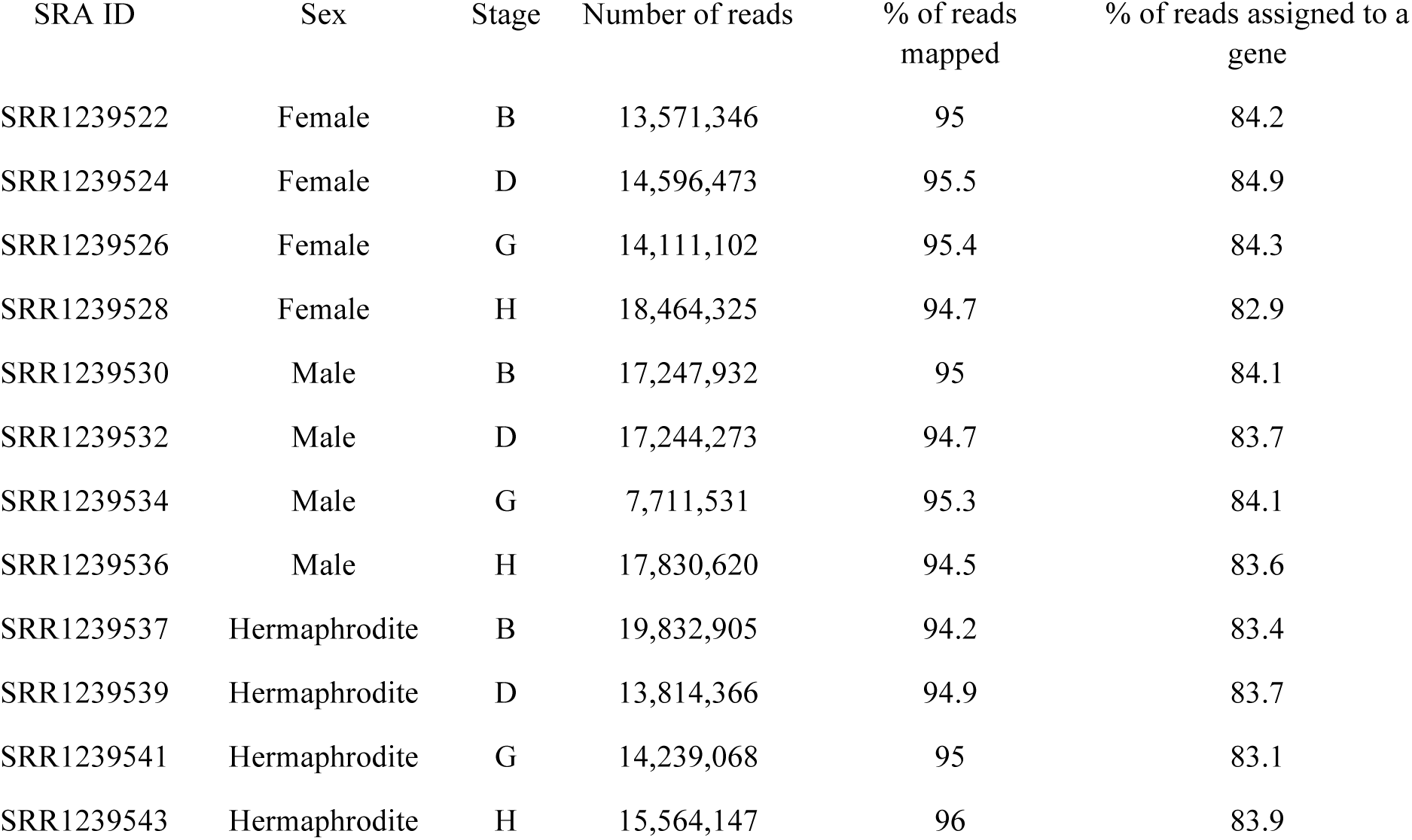
Mapping statistics of public RNA-seq libraries of flower buds of female, male and hermaphrodite *V. sylvestris* (Ramos et al. 2013) against the *V. sylvestris* genome. B to D represent successive developmental stages.

**Supplementary Table 12:**
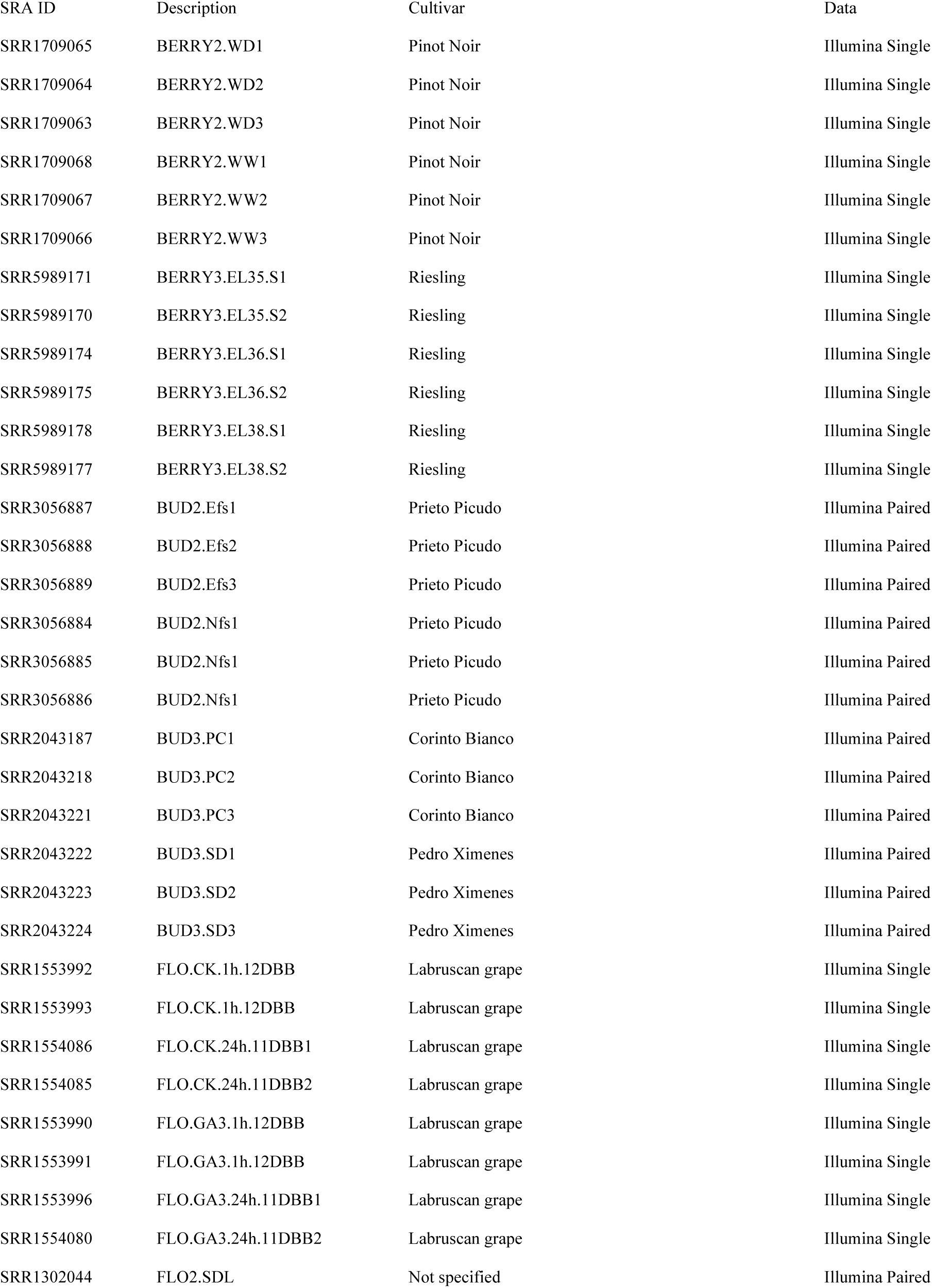
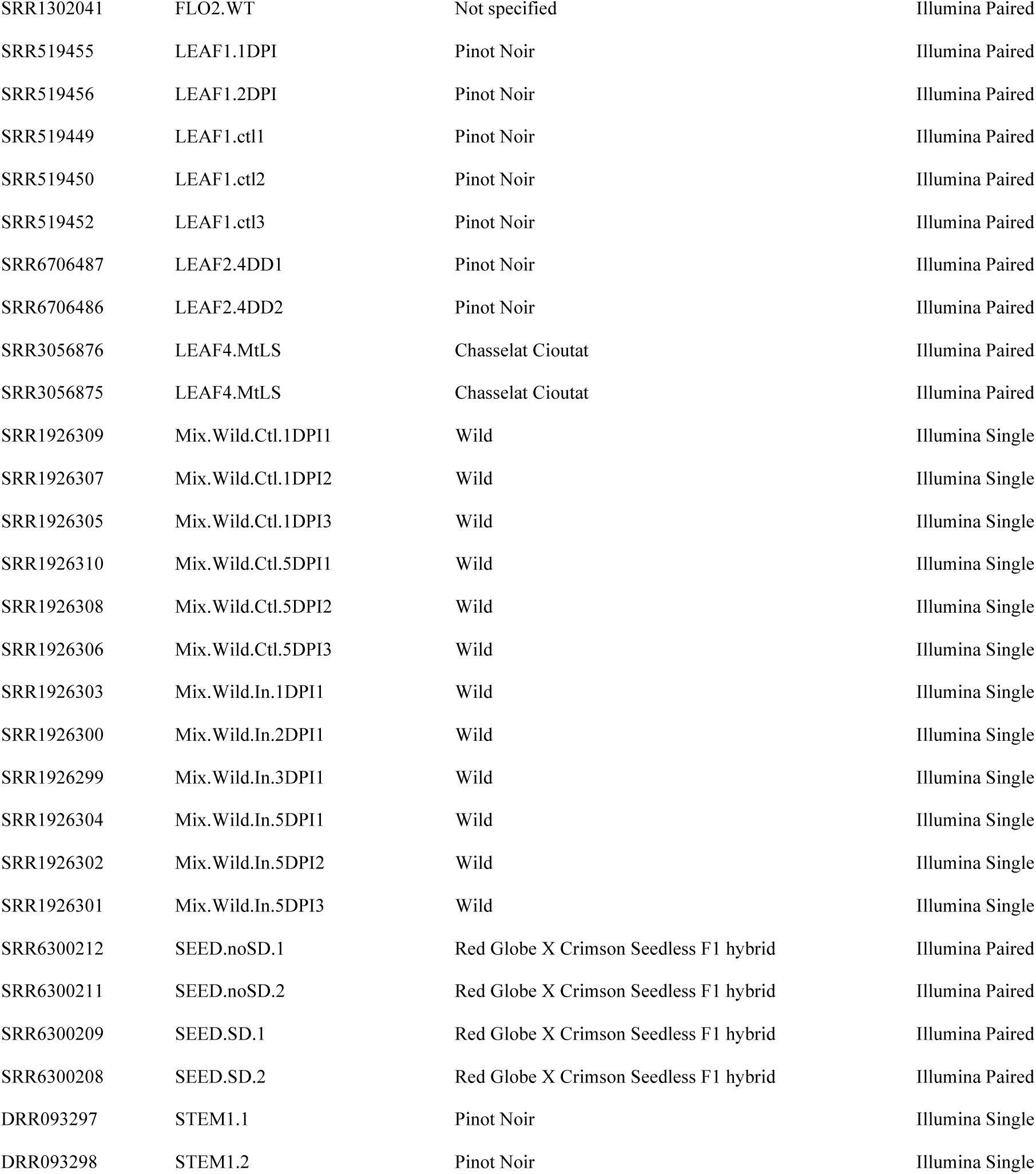
Public RNA-seq dataset of *V. vinifera* and *V. sylvestris* mapped to the *Vitis sylvestris* genome.

| SRA ID | Description | Cultivar | Data |
| --- | --- | --- | --- |
| SRR1709065 | BERRY2.WD1 | Pinot Noir | Illumina Single |
| SRR1709064 | BERRY2.WD2 | Pinot Noir | Illumina Single |
| SRR1709063 | BERRY2.WD3 | Pinot Noir | Illumina Single |
| SRR1709068 | BERRY2.WW1 | Pinot Noir | Illumina Single |
| SRR1709067 | BERRY2.WW2 | Pinot Noir | Illumina Single |
| SRR1709066 | BERRY2.WW3 | Pinot Noir | Illumina Single |
| SRR5989171 | BERRY3.EL35.S1 | Riesling | Illumina Single |
| SRR5989170 | BERRY3.EL35.S2 | Riesling | Illumina Single |
| SRR5989174 | BERRY3.EL36.S1 | Riesling | Illumina Single |
| SRR5989175 | BERRY3.EL36.S2 | Riesling | Illumina Single |
| SRR5989178 | BERRY3.EL38.S1 | Riesling | Illumina Single |
| SRR5989177 | BERRY3.EL38.S2 | Riesling | Illumina Single |
| SRR3056887 | BUD2.Efs1 | Prieto Picudo | Illumina Paired |
| SRR3056888 | BUD2.Efs2 | Prieto Picudo | Illumina Paired |
| SRR3056889 | BUD2.Efs3 | Prieto Picudo | Illumina Paired |
| SRR3056884 | BUD2.Nfs1 | Prieto Picudo | Illumina Paired |
| SRR3056885 | BUD2.Nfs1 | Prieto Picudo | Illumina Paired |
| SRR3056886 | BUD2.Nfs1 | Prieto Picudo | Illumina Paired |
| SRR2043187 | BUD3.PC1 | Corinto Bianco | Illumina Paired |
| SRR2043218 | BUD3.PC2 | Corinto Bianco | Illumina Paired |
| SRR2043221 | BUD3.PC3 | Corinto Bianco | Illumina Paired |
| SRR2043222 | BUD3.SD1 | Pedro Ximenes | Illumina Paired |
| SRR2043223 | BUD3.SD2 | Pedro Ximenes | Illumina Paired |
| SRR2043224 | BUD3.SD3 | Pedro Ximenes | Illumina Paired |
| SRR1553992 | FLO.CK.1h.12DBB | Labruscan grape | Illumina Single |
| SRR1553993 | FLO.CK.1h.12DBB | Labruscan grape | Illumina Single |
| SRR1554086 | FLO.CK.24h.11DBB1 | Labruscan grape | Illumina Single |
| SRR1554085 | FLO.CK.24h.11DBB2 | Labruscan grape | Illumina Single |
| SRR1553990 | FLO.GA3.1h.12DBB | Labruscan grape | Illumina Single |
| SRR1553991 | FLO.GA3.1h.12DBB | Labruscan grape | Illumina Single |
| SRR1553996 | FLO.GA3.24h.11DBB1 | Labruscan grape | Illumina Single |
| SRR1554080 | FLO.GA3.24h.11DBB2 | Labruscan grape | Illumina Single |
| SRR1302044 | FLO2.SDL | Not specified | Illumina Paired |
| SRR1302041 | FLO2.WT | Not specified | Illumina Paired |
| SRR519455 | LEAF1.1DPI | Pinot Noir | Illumina Paired |
| SRR519456 | LEAF1.2DPI | Pinot Noir | Illumina Paired |
| SRR519449 | LEAF1.ctl1 | Pinot Noir | Illumina Paired |
| SRR519450 | LEAF1.ctl2 | Pinot Noir | Illumina Paired |
| SRR519452 | LEAF1.ctl3 | Pinot Noir | Illumina Paired |
| SRR6706487 | LEAF2.4DD1 | Pinot Noir | Illumina Paired |
| SRR6706486 | LEAF2.4DD2 | Pinot Noir | Illumina Paired |
| SRR3056876 | LEAF4.MtLS | Chasselat Cioutat | Illumina Paired |
| SRR3056875 | LEAF4.MtLS | Chasselat Cioutat | Illumina Paired |
| SRR1926309 | Mix.Wild.Ctl.1DPI1 | Wild | Illumina Single |
| SRR1926307 | Mix.Wild.Ctl.1DPI2 | Wild | Illumina Single |
| SRR1926305 | Mix.Wild.Ctl.1DPI3 | Wild | Illumina Single |
| SRR1926310 | Mix.Wild.Ctl.5DPI1 | Wild | Illumina Single |
| SRR1926308 | Mix.Wild.Ctl.5DPI2 | Wild | Illumina Single |
| SRR1926306 | Mix.Wild.Ctl.5DPI3 | Wild | Illumina Single |
| SRR1926303 | Mix.Wild.In.1DPI1 | Wild | Illumina Single |
| SRR1926300 | Mix.Wild.In.2DPI1 | Wild | Illumina Single |
| SRR1926299 | Mix.Wild.In.3DPI1 | Wild | Illumina Single |
| SRR1926304 | Mix.Wild.In.5DPI1 | Wild | Illumina Single |
| SRR1926302 | Mix.Wild.In.5DPI2 | Wild | Illumina Single |
| SRR1926301 | Mix.Wild.In.5DPI3 | Wild | Illumina Single |
| SRR6300212 | SEED.noSD.1 | Red Globe X Crimson Seedless F1 hybrid | Illumina Paired |
| SRR6300211 | SEED.noSD.2 | Red Globe X Crimson Seedless F1 hybrid | Illumina Paired |
| SRR6300209 | SEED.SD.1 | Red Globe X Crimson Seedless F1 hybrid | Illumina Paired |
| SRR6300208 | SEED.SD.2 | Red Globe X Crimson Seedless F1 hybrid | Illumina Paired |
| DRR093297 | STEM1.1 | Pinot Noir | Illumina Single |
| DRR093298 | STEM1.2 | Pinot Noir | Illumina Single |

## Supplementary Text

### Results of BAC sequencing and assembly

The final results were: 1) a female haplotype composed of the overlapping of 08J18 and 20H04 BAC sequences, of a total 204,082 bp length. This haplotype maps correctly between position 4,866,319 and 5,119,669 of the chromosome 2 of the *Vitis vinifera* reference genome (cultivar PN40024, 12X.V0, NCBI), which is known to correspond to the female haplotype of the Pinot Noir cultivar (Fechter et al 2012, Picq et al 2014); and 2) a male haplotype composed of two non-overlapping contigs, contig1: BACs 27H5 and 65K18 (171,096bp) and contig2: BACs 9D16, 15F19 and 71N3 (169,751bp). The male haplotype does not map well on the grape genome reference, as the hermaphrodite haplotype of the PN40024 was not assembled on the chromosome 2 but its differing sequences were left unassigned (chromosome unknown). On the reference genome 12X.V0, the borders of the contig1 map at positions chr2:4,856,490 and chrUn:16,079,507; the borders of contig2 map at chr2:4,953,113 and chr:5,142,793. Based on the closest blast hits of the male contigs on the female and hermaphrodite haplotypes of the reference genome, as well as on the other available long- reads PacBio genomes (the Chardonnay sequences of Zhou et al 2019^19^; the Cabernet Sauvignon sequences of Chin et al 2016^9^; as well as the *Vitis sylvestris* female sequences of this work), we estimated the length of the gap between contig1 and contig2 at around 13kb.

### Covering the 13kb gap on the male haplotype

Further efforts were attempted to cover the gap between the two non-overlapping male contigs. As a gap of 13kb is too long to be covered by long-range PCR even when using high- quality TAQ, we made the hypothesis that the *PLATZ* gene, for which we have positional evidence in several female and hermaphrodite genotypes, is positioned within this gap in the male haplotype too, and can be used to split the 13kb in two shorter parts, easier to amplify by long-range PCR. The positions of *PLATZ* on H and F haplotypes were obtained from: the reference genome of PN40024, on which the *PLATZ* female allele maps at chr2:4949289..4950582 and its hermaphrodite allele at chrUn16091133..16092418; the Chardonnay PacBio sequences (Zhou et al 2019); the Cabernet Sauvignon PacBio sequences (Chin et al 2016); as well as the *Vitis sylvestris* female PacBio sequences (this work). In addition, the *PLATZ* male allele was also found by microassembly of reads carrying Y alleles detecting in this work from the males of the *Vitis sylvestris* cross (this work); for facility we call this male version *PLATZy*.

Based on this hypothetical construct (contig1-gap1-PLATZy-gap2-contig2), we defined several pairs of primers based on the contig1- and contig2 ends and on *PLATZy*. The primers were defined exploiting polymorphisms between the F and M haplotype, so to be sure to amplify only the M haplotype. Starting from primers based on *PLATZy* and contig2 (forward: ACTCCCCTGTTTCTCTCCGA and reverse: TCATGTTGCGTCTAGATCGGT), we were able to obtain a single band of 3.2kb, corresponding to gap2, by long-range PCR on the *Vitis sylvestris* PSL10 male. DNAs from one female *V. sylvestris* and the hermaphrodite Pinot noir (respectively with introduction codes 8500Mtp110 and 193Mtp81, INRA Vassal Grape Collection) did not provide any amplification, as expected. The forward and reverse Sanger sequences of this PCR product were obtained through an AB1 sequencer, and have 93-99% identity with the same region on 3’ of *PLATZ*, of the reference genome PN40024, and the Cabernet Sauvignon, the Chardonnay and the *Vitis sylvestris* female PacBio sequences. We can thus confirm that in the male haplotype, *PLATZy* is located in the expected region between *ETOL1* and the first *FMO*.

On the other hand, we did not succeed to sequence gap1, as PCR amplifications using several combinations of primers defined in contig1 and *PLATZ* always provided multiple bands, confirming that gap1 is highly likely to correspond to a repeated element, as observed in the reference genome PN40024, Cabernet Sauvignon, Chardonnay and *Vitis sylvestris* female.

## References

1. Renner, S. S. The relative and absolute frequencies of angiosperm sexual systems: Dioecy, monoecy, gynodioecy, and an updated online database. Am. J. Bot. 101, 1588– 1596 (2014).

2. Gallardo, A., Ocete, R., López, M. Á., Lara, M. & Rivera, D. Assessment of pollen dimorphism in populations of Vitis vinifera L. subsp. sylvestris (Gmelin) Hegi in Spain. 4.

3. Caporali, E., Spada, A., Marziani, G., Failla, O. & Scienza, A. The arrest of development of abortive reproductive organs in the unisexual flower of Vitis vinifera ssp. silvestris. Sex. Plant Reprod. 15, 291–300 (2003).

4. Fechter, I. et al. Candidate genes within a 143 kb region of the flower sex locus in Vitis. Mol. Genet. Genomics 287, 247–259 (2012).

5. Picq, S. et al. A small XY chromosomal region explains sex determination in wild dioecious V. vinifera and the reversal to hermaphroditism in domesticated grapevines. BMC Plant Biol. 14, 229 (2014).

6. Coito, J. L. et al. VviAPRT3 and VviFSEX: Two Genes Involved in Sex Specification Able to Distinguish Different Flower Types in Vitis. Front. Plant Sci. 8, (2017).

7. Chin, C.-S. et al. Phased diploid genome assembly with single-molecule real-time sequencing. Nat. Methods 13, 1050–1054 (2016).

8. Canaguier, A. et al. A new version of the grapevine reference genome assembly (12X.v2) and of its annotation (VCost.v3). Genomics Data 14, 56–62 (2017).

9. Raymond, O. et al. The Rosa genome provides new insights into the domestication of modern roses. Nat. Genet. 50, 772–777 (2018).

10. Badouin, H. et al. The sunflower genome provides insights into oil metabolism, flowering and Asterid evolution. Nature 546, 148–152 (2017).

11. Prentout, D. et al. A high-throughput segregation analysis identifies the sex chromosomes of Cannabis sativa. bioRxiv 721324 (2019) doi:10.1101/721324.

12. Muyle, A. et al. SEX-DETector: A Probabilistic Approach to Study Sex Chromosomes in Non-Model Organisms. Genome Biol. Evol. 8, 2530–2543 (2016).

13. Ramos, M. J. N. et al. Flower development and sex specification in wild grapevine. BMC Genomics 15, 1095 (2014).

14. Zhou, Y., Massonnet, M., Sanjak, J. S., Cantu, D. & Gaut, B. S. Evolutionary genomics of grape (Vitis vinifera ssp. vinifera) domestication. Proc. Natl. Acad. Sci. 114, 11715– 11720 (2017).

15. Veltsos, P. et al. Size and Content of the Sex-Determining Region of the Y Chromosome in Dioecious Mercurialis annua, a Plant with Homomorphic Sex Chromosomes. Genes 9, (2018).

16. Charlesworth, B. The evolution of sex chromosomes. Science 251, 1030–1033 (1991).

17. Zhou, Y. et al. The population genetics of structural variants in grapevine domestication. Nat. Plants 5, 965–979 (2019).

18. Dobritsa, A. A. & Coerper, D. The novel plant protein INAPERTURATE POLLEN1 marks distinct cellular domains and controls formation of apertures in the Arabidopsis pollen exine. Plant Cell 24, 4452–4464 (2012).

19. Dobritsa, A. A., Kirkpatrick, A. B., Reeder, S. H., Li, P. & Owen, H. A. Pollen Aperture Factor INP1 Acts Late in Aperture Formation by Excluding Specific Membrane Domains from Exine Deposition. Plant Physiol. 176, 326–339 (2018).

20. Furness, C. A. Why does some pollen lack apertures? A review of inaperturate pollen in eudicots. Bot. J. Linn. Soc. 155, 29–48 (2007).

21. Golovkin, M. & Reddy, A. S. N. A calmodulin-binding protein from Arabidopsis has an essential role in pollen germination. Proc. Natl. Acad. Sci. 100, 10558–10563 (2003).

22. Boualem, A. et al. A Conserved Ethylene Biosynthesis Enzyme Leads to Andromonoecy in Two Cucumis Species. PLOS ONE 4, e6144 (2009).

23. Li, X. et al. Rice APOPTOSIS INHIBITOR5 Coupled with Two DEAD-Box Adenosine 5′-Triphosphate-Dependent RNA Helicases Regulates Tapetum Degeneration. Plant Cell 23, 1416–1434 (2011).

24. Akagi, T. et al. A Y-Encoded Suppressor of Feminization Arose via Lineage-Specific Duplication of a Cytokinin Response Regulator in Kiwifruit. Plant Cell 30, 780–795 (2018).

25. Akagi, T. et al. Two Y-chromosome-encoded genes determine sex in kiwifruit. Nat. Plants 5, 801–809 (2019).

26. Akagi, T., Henry, I. M., Tao, R. & Comai, L. A Y-chromosome-encoded small RNA acts as a sex determinant in persimmons. Science 346, 646–650 (2014).

27. Negi, S. S. & Olmo, H. P. Sex Conversion in a Male Vitis vinifera L. by a Kinin. Science 152, 1624–1624 (1966).

28. Käfer, J., Marais, G. A. B. & Pannell, J. R. On the rarity of dioecy in flowering plants. Mol. Ecol. 26, 1225–1241 (2017).

## Supplementary References

29. R Core Team. R: A Language and Environment for Statistical Computing. (R Foundation for Statistical Computing, 2014).

30. Krzywinski, M. et al. Circos: An information aesthetic for comparative genomics. Genome Res. 19, 1639–1645 (2009).

31. Jombart, T. adegenet: a R package for the multivariate analysis of genetic markers. Bioinformatics 24, 1403–1405 (2008).

32. Li, H. & Durbin, R. Fast and accurate long-read alignment with Burrows-Wheeler transform. Bioinformatics 26, 589–595 (2010).

33. Simao, F. A., Waterhouse, R. M., Ioannidis, P., Kriventseva, E. V. & Zdobnov, E. M. BUSCO: Assessing genome assembly and annotation completeness with single-copy orthologs. Bioinformatics 31, 3210–3212 (2015).

34. Roach, M. J. et al. Population sequencing reveals clonal diversity and ancestral inbreeding in the grapevine cultivar Chardonnay. PLOS Genet. 14, e1007807 (2018).

35. Peterson, D. G., Tomkins, J. P., Frisch, D. A., Wing, A. & Paterson, A. H. Construction of Plant Bacterial Artificial Chromosome (BAC) Libraries: An Illustrated Guide. 2nd Edition. J. Agric. Genomics 5, (2000).

36. Schweiger, W. et al. Suppressed recombination and unique candidate genes in the divergent haplotype encoding Fhb1, a major Fusarium head blight resistance locus in wheat. Theor. Appl. Genet. 129, 1607–1623 (2016).

37. Schmieder, R. & Edwards, R. Quality control and preprocessing of metagenomic datasets. Bioinformatics 27, 863–864 (2011).

38. Schmieder, R., Lim, Y. W. & Edwards, R. Identification and removal of ribosomal RNA sequences from metatranscriptomes. Bioinformatics 28, 433–435 (2012).

39. Dobin, A. et al. STAR: ultrafast universal RNA-seq aligner. Bioinformatics 29, 15–21 (2013).

40. Grabherr, M. G. et al. Full-length transcriptome assembly from RNA-Seq data without a reference genome. Nat. Biotechnol. 29, 644–52 (2011).

41. Sallet, E., Gouzy, J. & Schiex, T. EuGene: An Automated Integrative Gene Finder for Eukaryotes and Prokaryotes. Methods Mol. Biol. Clifton NJ 1962, 97–120 (2019).

42. Wu, T. D. & Watanabe, C. K. GMAP: a genomic mapping and alignment program for mRNA and EST sequences. Bioinformatics 21, 1859–1875 (2005).

43. Li, H. et al. The Sequence Alignment/Map format and SAMtools. Bioinformatics 25, 2078–2079 (2009).

44. Koboldt, D. C. et al. VarScan 2: Somatic mutation and copy number alteration discovery in cancer by exome sequencing. Genome Res. 22, 568–576 (2012).

45. Yang, Z. PAML 4: Phylogenetic Analysis by Maximum Likelihood. Mol. Biol. Evol. 24, 1586–1591 (2007).

46. Girgis, H. Z. Red: an intelligent, rapid, accurate tool for detecting repeats de-novo on the genomic scale. BMC Bioinformatics 16, 227 (2015).

47. Katoh, K., Rozewicki, J. & Yamada, K. D. MAFFT online service: multiple sequence alignment, interactive sequence choice and visualization. Brief. Bioinform. 20, 1160–1166 (2019).

48. Anders, S., Pyl, P. T. & Huber, W. HTSeq—a Python framework to work with high- throughput sequencing data. Bioinformatics 31, 166–169 (2015).

49. Quinlan, A. R. & Hall, I. M. BEDTools: a flexible suite of utilities for comparing genomic features. Bioinformatics 26, 841–842 (2010).

50. Yanai, I. et al. Genome-wide midrange transcription profiles reveal expression level relationships in human tissue specification. Bioinformatics 21, 650–659 (2005).

